# Secreted phospholipase PLA2G12A-driven lysophospholipid signaling via lipolytic modification of extracellular vesicles facilitates pathogenic Th17 differentiation

**DOI:** 10.1101/2024.10.27.620543

**Authors:** Chika Mochizuki, Yoshitaka Taketomi, Atsushi Irie, Kuniyuki Kano, Yuki Nagasaki, Yoshimi Miki, Takashi Ono, Yasumasa Nishito, Takahiro Nakajima, Yuri Tomabechi, Kazuharu Hanada, Mikako Shirouzu, Takashi Watanabe, Kosuke Hata, Yoshihiro Izumi, Takeshi Bamba, Jerold Chun, Kai Kudo, Ai Kotani, Yusuke Endo, Junken Aoki, Makoto Murakami

**Author notes:** These authors equally contributed to this work (Chika Mochizuki, Yoshitaka Taketomi).

## Abstract

Lipogenesis-driven metabolic flux is crucial for differentiation of pathogenic Th17 cells. Although our previous CRISPR-based screening identified PLA2G12A as a key player in this process, it has remained obscure how this secreted phospholipase A_2_ isoform controls Th17 differentiation. Here we show that global, T cell-specific, or fibroblast-specific deletion of PLA2G12A prevents Th17 differentiation and associated diseases including psoriasis and arthritis. PLA2G12A acts on Th17-derived extracellular vesicles (EVs) to produce lysophospholipids including the RORγt activator 1-oleoyl-lysophosphatidylethanolamine. These lysophospholipids are further converted by autotaxin to lysophosphatidic acid (LPA), which assists Th17 differentiation mainly via LPA_2_ receptor. Moreover, PLA2G12A promotes the secretion and uptake of EVs by Th17 cells and alters their cargo contents. Defective Th17 differentiation by PLA2G12A deficiency is rescued by supplementation with PLA2G12A-modified EVs. Importantly, a PLA2G12A-blocking antibody prevents Th17 differentiation and ameliorates psoriasis and arthritis models. Thus, targeting the PLA2G12A-EV-lysophospholipid axis may be useful for treatment of Th17-related diseases.

## INTRODUCTION

T helper 17 (Th17) cells, which differentiate from naive T cells primarily through the combined action of IL-6 and TGF-β, contribute to host defense against extracellular bacteria and fungi at epithelial surfaces ^1^. Th17 cytokines including IL-17A, IL-17F and IL-22, which are also produced by innate-type lymphocytes such as γδ T cells and type 3 innate lymphoid cells (ILC3), play crucial roles in maintenance of tissue homeostasis by inducing antimicrobial peptides and protecting the epithelial barrier against pathogens ^2-4^. IL-23 and IL-1β supplied by dendritic cells (DCs) or macrophages in tissue microenvironments increase the pathogenicity of Th17 cells in various autoimmune diseases such as psoriasis, rheumatoid arthritis, multiple sclerosis, inflammatory bowel disease, and type 2 diabetes ^5-8^. Homeostatic and pathogenic Th17 cells have distinct gene signatures, creating Th17 heterogeneity ^9-11^. Identification of the specific molecular switches that drive pathogenic *versus* homeostatic Th17 cells would allow selective inhibition of pathogenic Th17 cells while sparing non-pathogenic, tissue-protective Th17 cells.

Lipid metabolism crucially influences the differentiation and function of Th17 cells ^12-16^. Intake of fat-rich diet exacerbates Th17-related diseases such as psoriasis and type 2 diabetes ^17,18^. RORγt, the master transcription factor for Th17 cells, requires the binding of lipophilic ligands such as vitamins, steroids, retinoids, and fatty acids for its transactivation ^19^. Oxysterols or cholesterol-biosynthetic intermediates (*e.g*., lanosterol) have the capacity to increase RORγt activity in Th17 cells ^20,21^ and to activate γδ T cells via GPR183 ^22^. Loss of CD5L converts homeostatic Th17 cells to pathogenic Th17 cells by altering the lipid profile, with a decrease in cholesterol biosynthesis as well as an increase in saturated fatty acids (SFAs) over polyunsaturated fatty acids (PUFAs), eventually modulating the ligand availability to RORγt ^23^. Palmitoylation of STAT3 by the palmitoyltransferase DHHC7 favors its membrane recruitment and activation, thereby augmenting Th17 differentiation ^24^. Moreover, T cell-intrinsic signaling of prostaglandin E_2_ (PGE_2_), an arachidonic acid (AA; ω6 20:4) metabolite, via its receptor EP4 is involved in Th17-driven inflammation ^25^, exemplifying the regulation of pathogenic Th17 cells by a specific lipid mediator.

Pharmacological inhibition or genetic ablation of acetyl-CoA carboxylase (ACC1) and fatty acid synthase (FASN), which catalyze the first and second steps of fatty acid synthesis, respectively, in T cells impairs Th17 differentiation and increases regulatory T (T_reg_) cells, which can be reversed by supplementation with exogenous fatty acids ^26-28^, suggesting that fatty acid metabolism dictates Th17 and T_reg_ balance. By exploiting the advantages of CRISPR-based screening along with unbiased lipidomic analysis, we have recently shown that several lipogenic enzymes, including SCD2 (fatty acid desaturase), GPAM, GPAM3 and LPLAT1 (enzymes involved in glycerophospholipid (hereafter phospholipid) synthesis), operate downstream of ACC1 and FASN to increase the transcriptional activity of RORγt for a core molecular signature of Th17 cells and that 1-oleoyl-lysophosphatidylethanolamine [LPE(1-18:1)], a lysophospholipid with *sn*-1 oleic acid (OA; 18:1), a monounsaturated fatty acid (MUFA), serves as an endogenous ligand for RORγt ^29^. However, behind MUFA and phospholipid generation, the precise biosynthetic route for LPE(1-18:1) during Th17 differentiation has not yet been fully clarified.

Lysophospholipids are produced primarily by hydrolysis of phospholipids by phospholipase A_2_ (PLA_2_) or A_1_ (PLA_1_) enzymes. The mammalian genome encodes more than 50 PLA_2_s or related enzymes, which are subdivided into several families including cytosolic PLA_2_s (cPLA_2_s), Ca^2+^-independent PLA_2_s (iPLA_2_s), secreted PLA_2_s (sPLA_2_s), and others ^30^. Besides the aforementioned lipogenic enzymes, CRISPR-based deletion of PLA2G12A (sPLA_2_-XIIA), an atypical member of the sPLA_2_ family with an unknown function ^31,32^, in T cells has been shown to hamper Th17 differentiation ^29^, suggesting that this sPLA_2_ is a candidate enzyme responsible for the generation of LPE(1-18:1). However, it remains unresolved whether PLA2G12A indeed generates LPE(1-18:1) and/or additional lipid metabolite(s) potentially acting in concert with LPE(1-18:1) for Th17 differentiation, and if so, which membrane serves as the hydrolytic target of this sPLA_2_, how and where its genetic deletion affects Th17-related diseases, and whether the PLA2G12A-driven lipid pathway would be a druggable target.

In the present study, we have provided evidence that PLA2G12A does indeed act as a critical regulator of Th17 differentiation by hydrolyzing Th17-derived EVs to produce LPE(1-18:1) and secondarily lysophosphatidic acid (LPA), which acts on G_12/13_- coupled LPA_2_ and possibly LPA_1_ receptors to promote Th17 differentiation. PLA2G12A also alters the properties of Th17-derived EVs and modifies their secretion, uptake, and cargo contents. Importantly, antibody-mediated blocking of PLA2G12A markedly attenuates mouse models of psoriasis and arthritis by preventing Th17 differentiation, implicating PLA2G12A as a novel target for drug treatment of Th17-driven diseases.

## RESULTS

### PLA2G12A deficiency impairs Th17 differentiation *ex vivo*

Quantitative RT-PCR (qPCR) revealed that *Pla2g12a* mRNA was ubiquitously expressed in C57BL/6 mouse tissues, with relatively high expression in the lung, stomach, white (WAT) and brown (BAT) adipose tissues, and lymphoid tissues including the spleen and lymph nodes (LNs) (Figure S1A). A similar expression profile of *PLA2G12A* was also seen in humans (https://www.proteinatlas.org/ENSG00000123739-PLA2G12A/tissue). To assess the expression of *Pla2g12a* in Th17 cells, CD4^+^CD62L^+^ naïve T cells isolated from mouse spleen were cultured for 3 days in the presence (for Th17) or absence (for Th0) of a cytokine cocktail containing IL-1β, IL-6, IL-23 and TGF-β and an anti-CD28 antibody in culture plates precoated with an anti-CD3ε antibody (Figures 1A and S1B). In this *ex vivo* T cell culture, *Pla2g12a* expression was markedly increased in Th0 cells and even further expressed in Th17 cells after 6-24 h, followed by a decline after 48 h of culture (Figure 1B), suggesting that its expression is transiently induced by T cell receptor (TCR) activation and augmented by Th17 polarization. Among the various PLA_2_ enzymes (sPLA_2_s, cPLA_2_s and iPLA_2_s), *Pla2g12a* was the most highly expressed PLA_2_ in Th0 cells and the only PLA_2_ that was further increased in Th17 cells (Figure S1C).

**Figure 1.**
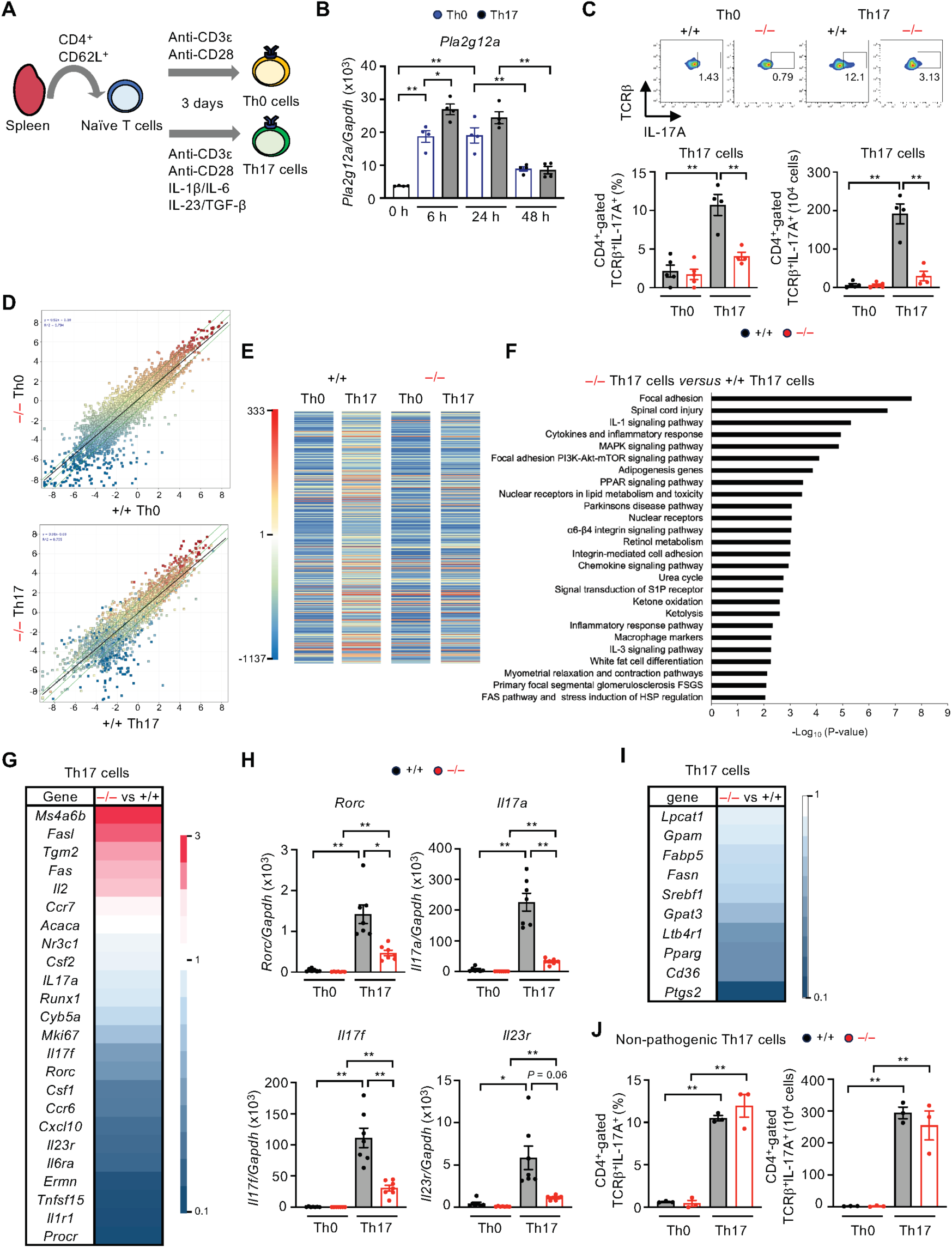
Impaired *ex vivo* Th17 differentiation in *Pla2g12a*^-/-^ mice. (A) Schematic representation of the procedure for *ex vivo* culture of splenic naïve CD4^+^ T cells with (Th17) or without (Th0) IL-1β, IL-6, IL-23 and TGF-β on anti-CD3ε/CD28-coated plates. (B) qPCR of *Pla2g12a* in splenic CD4^+^ T cells cultured for various periods under Th0 and Th17 differentiation conditions. (C) FACS of Th17 (CD4^+^-gated TCRβ^+^IL-17A^+^) cells after culture of naïve T cells prepared from *Pla2g12a*^+/+^ (+/+) and *Pla2g12a*^-/-^ (–/–) mice for 3 days under Th0 and Th17 differentiation conditions. Representative FACS profiles (*left*) and the proportion and number of Th17 cells (*right*) are shown. (D-F) Scatter plots (D) and heatmap (E) of genes expressed in CD4^+^ T cells from *Pla2g12a*^+/+^ and *Pla2g12a*^-/-^ mice after culture for 3 days under Th0 and Th17 differentiation conditions. Cells from 6-7 mice of each genotype were pooled and subjected to microarray analysis. Colors in (E) show the z-score. (F) Pathway enrichment analysis of genes decreased (>2-fold) in *Pla2g12a*^-/-^ Th17 cells relative to *Pla2g12a*^+/+^ Th17 cells. (G) Heatmap of Th17-related genes in Th17 cells from *Pla2g12a*^-/-^ mice relative to those from *Pla2g12a*^+/+^ mice. Colors indicate fold changes in *Pla2g12a*^-/-^ cells relative to *Pla2g12a*^+/+^ cells. (H) qPCR of Th17 signature genes in Th0 and Th17 cells from *Pla2g12a*^+/+^ and *Pla2g12a*^-/-^ mice. (I) Heatmap of lipid-related genes in *Pla2g12a*^-/-^ Th17 cells relative to *Pla2g12a*^+/+^ Th17 cells. Colors indicate fold changes in *Pla2g12a*^-/-^ cells relative to *Pla2g12a*^+/+^ cells. (J) FACS of non-pathogenic Th17 cells from *Pla2g12a*^+/+^ and *Pla2g12a*^-/-^ mice after culture for 3 days with IL-6 and TGF-β. Values are mean ± s.e.m.. Representative data of three experiments (C) and results of one experiment (B, D–I) are shown. Statistical analysis was performed by one way ANOVA with Tukey’s multiple comparisons test (C, J), Brown-Forsythe and Welch ANOVA with Dunnett’s T3 multiple comparisons test (H), and two-way ANOVA with Sidak’s multiple comparisons test (B). *, *P* < 0.05; **, *P* < 0.01.

To address the *in vivo* roles of PLA2G12A, we generated *Pla2g12a*^-/-^ mice, lacking PLA2G12A in the whole body, on a C57BL/6 genetic background (Figures S1D and S1E). qPCR confirmed that *Pla2g12a* mRNA was undetectable in the skin and spleen of *Pla2g12a*^-/-^ mice (Figure S1F). Under normal housing conditions, *Pla2g12a*^-/-^ mice were born normally and fertile, and developed no gross abnormalities beyond 1 year of age. In *ex vivo* T cell culture, Th17 differentiation (as monitored by flow cytometry of CD4-gated TCRβ^+^IL-17A^+^ cells) of naïve T cells obtained from *Pla2g12a*^-/-^ mice was markedly impaired as compared with those from *Pla2g12a*^+/+^ mice (Figure 1C). Microarray gene profiling revealed a marked difference in gene expression profiles between *Pla2g12a*^+/+^ and *Pla2g12a*^-/-^ Th17 cells (Figures 1D and 1E). Even in Th0 cells, a substantial set of genes were differentially expressed between the genotypes (Figure 1D), suggesting that PLA2G12A deficiency had some influence on CD4^+^ T cells even without Th17 differentiation, possibly following TCR stimulation. Nevertheless, heatmap visualization clearly indicated that numerous genes were dramatically upregulated in *Pla2g12a*^+/+^ Th17 cells relative to Th0 cells, whereas this event was only modest in *Pla2g12a*^-/-^ Th17 cells (Figure 1E). Gene ontology (GO) analysis showed that genes related to cytokine and chemokine signaling, cell proliferation and adhesion, inflammatory responses, and lipid metabolism were downregulated in *Pla2g12a*^-/-^ Th17 cells relative to *Pla2g12a*^+/+^ Th17 cells (Figure 1F). Decreased expression of genes related to the pathogenic Th17 program (*e.g., Rorc, Il17a, Csf2, Mki67, Il23r, Ermn, Il1r1*, and *Procr*) and increased expression of genes related to a non-pathogenic, stem-like nature (*e.g., Ms4a6b, Fasl, Tgm2, Fas,* and *Il2*) ^11^ were evident in *Pla2g12a*^-/-^ cells relative to *Pla2g12a*^+/+^ cells (Figure 1G). qPCR confirmed that loss of PLA2G12A impaired the induction of the core Th17-related genes *Rorc, Il17a, Il17f* and *Il23r* in Th17 cells (Figure 1H). Furthermore, expression levels of genes related to lipid synthesis, uptake, and signaling (*e.g*., *Srebf1*, *Pparg*, *Cd36*, *Fabp5, Ltb4r1* and *Ptgs2*) were noticeably reduced in *Pla2g12a*^-/-^ relative to *Pla2g12a*^+/+^ Th17 cells (Figure 1I). In contrast, PLA2G12A deletion affected the differentiation of non-pathogenic Th17 cells by IL-6 and TGF-β only minimally (Figure 1J). Thus, consistent with our previous study ^29^, PLA2G12A contributes to the differentiation of pathogenic Th17 cells *ex vivo*.

### PLA2G12A deficiency attenuates imiquimod (IMQ)-induced psoriasis

To assess the role of PLA2G12A in Th17-related pathology *in vivo*, we used IMQ-induced skin inflammation, a well-known model of human psoriasis vulgaris that depends on IL-17A-producing T cells ^33^, in *Pla2g12a*^+/+^ and *Pla2g12a*^-/-^ mice. The experimental procedure is illustrated in Figure 2A. *Pla2g12a* expression in the LNs was increased on day 1 and declined thereafter (Figure 2B), in agreement with its transient upregulation after TCR activation (Figure 1B). IMQ-elicited ear swelling over time (Figure 2C) and epidermal and dermal thickening on day 6 (Figures 2D and 2E) were significantly attenuated in *Pla2g12a*^-/-^ mice relative to *Pla2g12a*^+/+^ mice. IMQ-induced *Il17a* expression in the LNs and ear skin was lower in IMQ-treated *Pla2g12a*^-/-^ mice than in replicate *Pla2g12a*^+/+^ mice (Figure 2F). In contrast, expression levels of *Il6*, *Il22,* and *Il23p19* in the LNs and skin at each time point were similar in both genotypes (Figures S2A and S2B), suggesting that Th22 cells and DCs were not profoundly affected by loss of PLA2G12A. Of the keratinocyte markers *Krt10* and *Flg,* increased expression of the latter on day 6 was lower in *Pla2g12a*^-/-^ mice than in *Pla2g12a*^+/+^ mice (Figure S2C), possibly reflecting a secondary effect of the skin pathology. FACS analysis of the spleen and ear skin confirmed that the proportion and number of TCRβ^+^IL-17A^+^ Th17 cells on day 6 after IMQ challenge were significantly attenuated in *Pla2g12a*^-/-^ mice relative to *Pla2g12a*^+/+^ mice (Figures 2G, 2H, S2D and S2E). Since γδ T cells are one of the major sources of IL-17A in psoriatic skin ^34-36^, we also evaluated γδ T cells in ear skin and found that the increase of TCRγδ^+^IL-17A^+^ T cells after IMQ treatment was less obvious in *Pla2g12a*^-/-^ mice than in *Pla2g12a*^+/+^ mice (Figures 2I and S2E). Thus, PLA2G12A ablation selectively impairs the IMQ-induced increase in IL-17A-producing T cells including Th17 and γδ T cells.

**Figure 2.**
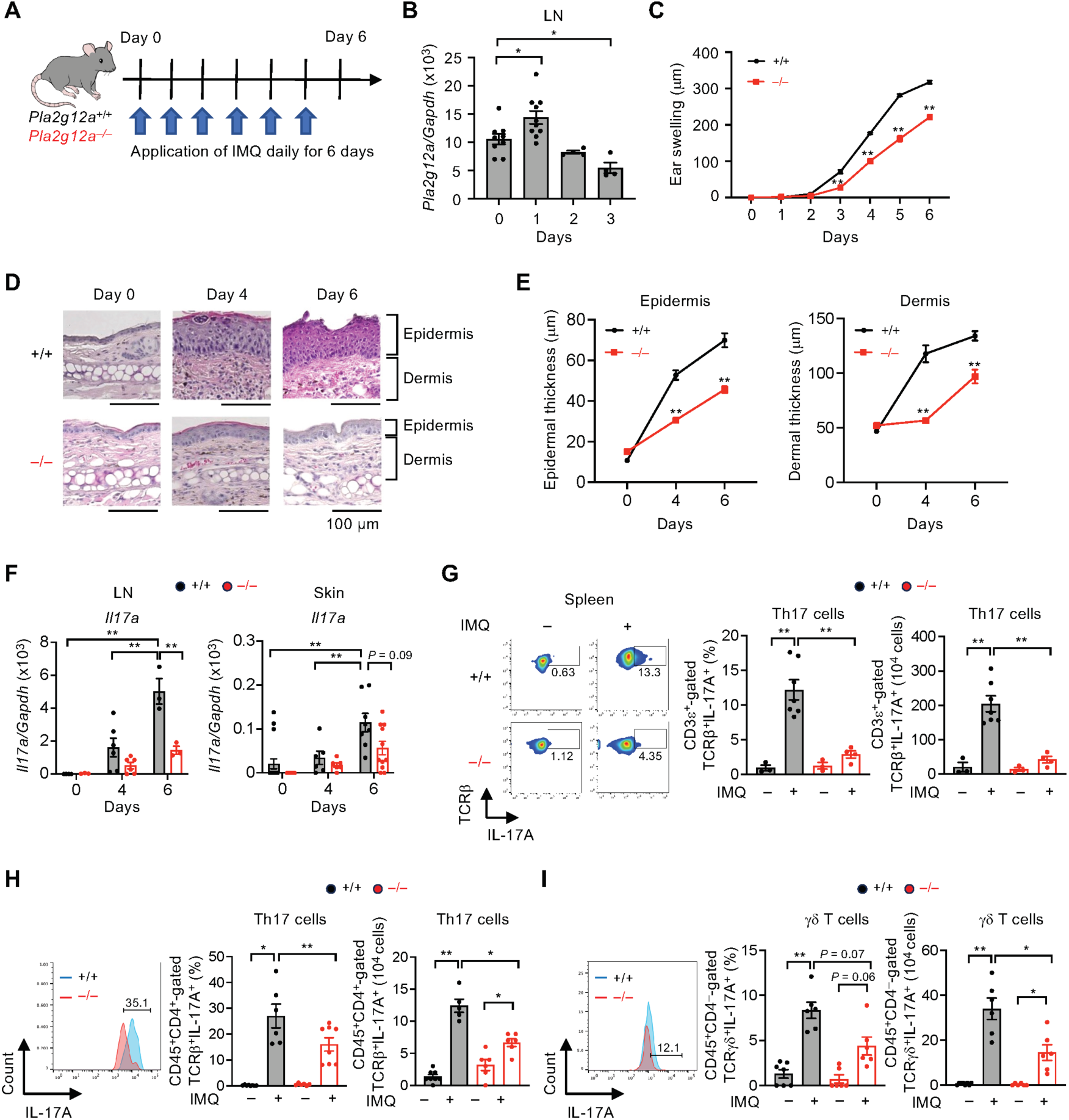
Amelioration of IMQ-induced psoriasis by global PLA2G12A deficiency. (A) Schematic representation of the procedure for IMQ-induced psoriasis. (B) qPCR of *Pla2g12a* in the LNs of WT mice after IMQ challenge over 3 days. (C) IMQ-induced ear swelling in *Pla2g12a*^+/+^ (+/+) and *Pla2g12a*^-/-^ (–/–) mice over 6 days (*n* = 6–18). (D, E) HE staining of ear skin sections (D) and quantification of epidermal and dermal thickness (E) in *Pla2g12a*^+/+^ and *Pla2g12a*^-/-^ mice (*n* = 35–44) (E). Scale bar, 100 μm. (F) qPCR of *Il17a* in the LNs and skin of *Pla2g12a*^+/+^ and *Pla2g12a*^-/-^ mice. (G) FACS of Th17 (CD3ε^+^-gated TCRβ^+^IL-17A^+^) cells in the spleen of *Pla2g12a*^+/+^ and *Pla2g12a*^-/-^ mice on day 6. Representative FACS profiles (*left*) and the proportion and number of Th17 cells (*right*) are shown. (H, I) FACS of Th17 (CD45^+^-gated TCRβ^+^IL-17A^+^) cells (H) and γδ T (CD45^+^-gated TCRγδ^+^IL-17A^+^) cells (I) in the skin on day 6. Representative FACS profiles (*left*) and the proportions and numbers of Th17 cells (H) and γδ T cells (I) (*right*) are shown. Values are mean ± s.e.m.. Representative data of two (C, F) or four (G) experiments, results from one experiment (H, I), and combined results of two experiments (B, D, E) are shown. Statistical analysis was performed using ordinary one-way ANOVA with Dunnett’s multiple comparisons test (B), one-way ANOVA with Tukey’s multiple comparisons test (G), Brown-Forsythe and Welch ANOVA with Dunnett’s T3 multiple comparisons test (H, I), Mixed-effects model with Sidak’s multiple comparisons test (C), and two-way ANOVA with Sidak’s multiple comparisons test (E, F). *, *P* < 0.05; **, *P* < 0.01.

To address whether PLA2G12A expressed in CD4^+^ T cells would indeed contribute to psoriasis-like pathology, *Pla2g12a*^fl/fl^ mice were crossed with transgenic mice overexpressing Cre recombinase under the *Cd4* promoter to obtain helper T cell-specific PLA2G12A-deficient (*Pla2g12a*^fl/fl^*Cd4*^cre^) mice ^29^. Although *Pla2g12a* expression in the spleen was reduced only modestly, it was reduced markedly in CD4^+^ T cells isolated from the spleen of *Pla2g12a*^fl/fl^*Cd4*^cre^ mice relative to control *Pla2g12a*^fl/fl^ mice (Figure 3A), thus confirming successful ablation of PLA2G12A in CD4^+^ T cells*. Ex vivo* Th17 differentiation of *Pla2g12a*^fl/fl^*Cd4*^cre^ naïve T cells was markedly impaired relative to that of *Pla2g12a*^fl/fl^ cells (Figure 3B). When these mice were challenged with IMQ *in vivo*, ear swelling (Figure 3C), epidermal thickening and dermal inflammatory cell infiltration (Figure 3D), the percentages of Th17 and γδ T cells in the spleen and skin (Figures 3E and 3F), and cutaneous *Il17a* expression (Figure 3G) were markedly lower in *Pla2g12a*^fl/fl^*Cd4*^cre^ mice than in *Pla2g12a*^fl/fl^ mice, thus verifying the critical contribution of PLA2G12A in CD4^+^ T cells to psoriasis-like pathology, even though it was expressed mainly in a CD4^-^ cell population within the spleen.

**Figure 3.**
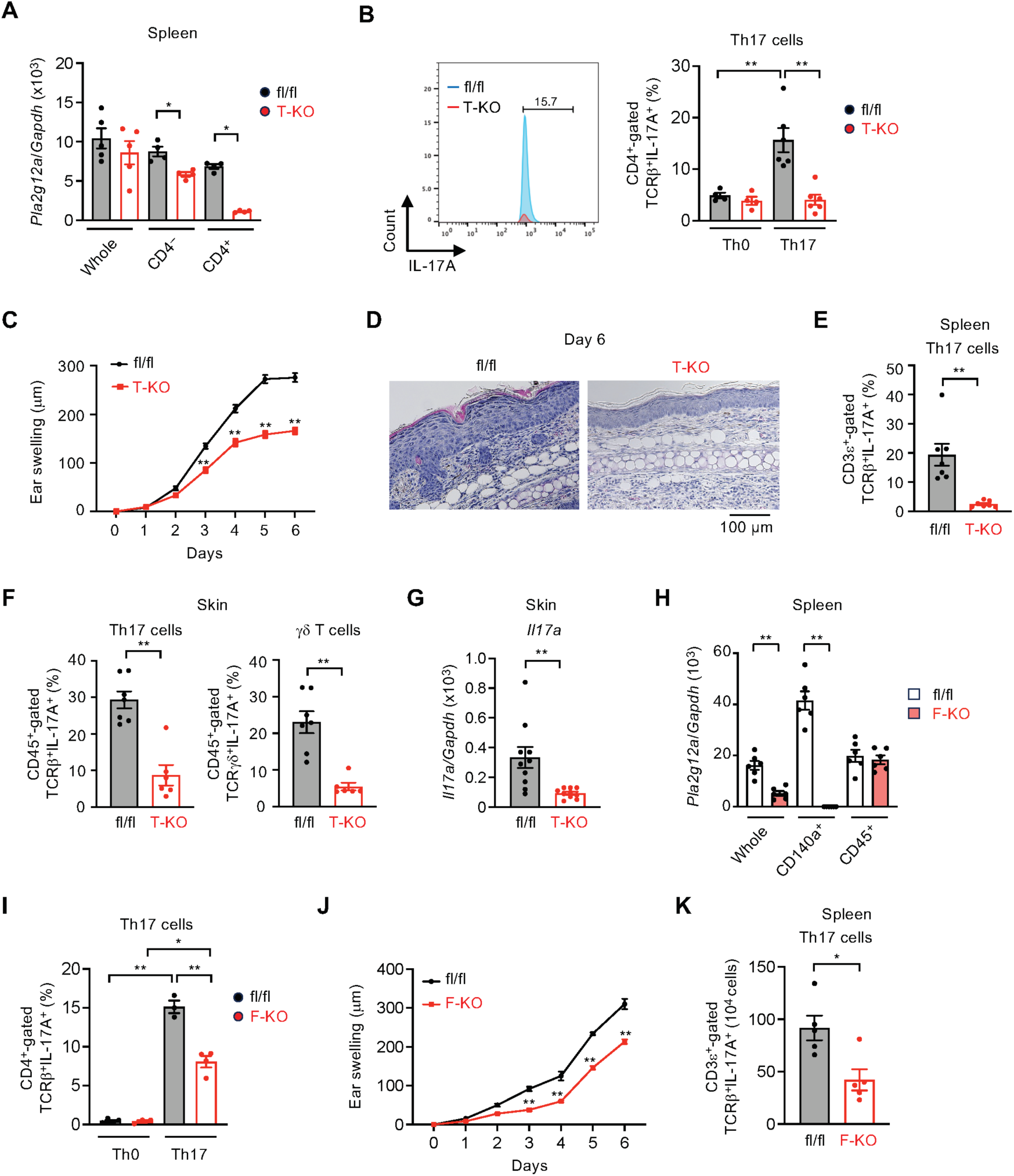
Amelioration of IMQ-induced psoriasis by CD4^+^ T cell- or fibroblast-specific PLA2G12A deficiency. (A) qPCR of *Pla2g12a* in the whole spleen and isolated CD4^-^ and CD4^+^ cells of *Pla2g12a*^fl/fl^ (fl/fl) and *Pla2g12a*^fl/fl^*Cd4*^cre^ (T-KO) mice. (B) FACS of Th17 cells after culture of naïve T cells from fl/fl and T-KO mice for 3 days under Th0 and Th17 differentiation conditions. Representative FACS profiles (*left*) and the proportion of Th17 cells (*right*) are shown. (C) IMQ-induced ear swelling in fl/fl and T-KO mice over 6 days (*n* = 20–30). (D) HE staining of ear skin sections from IMQ-treated fl/fl and T-KO mice on day 6. Scale bar, 100 μm. (E, F) FACS of Th17 cells in the spleen (E) and Th17 and γδ T cells in the skin (F) of IMQ-treated fl/fl and T-KO mice on day 6. (G) qPCR of *Il17a* in the skin of IMQ-treated fl/fl and T-KO mice on day 6. (H) qPCR of *Pla2g12a* in the whole LNs and isolated CD45^+^ hematopoietic cells and CD140a^+^ fibroblasts of fl/fl and *Pla2g12a*^fl/fl^*Col1a2*^cre^ (F-KO) mice. (I) FACS of Th17 cells after culture of naïve T cells from fl/fl and F-KO mice for 3 days under Th0 and Th17 differentiation conditions. Representative FACS profiles (*left*) and the proportion of Th17 cells (*right*) are shown. (J) IMQ-induced ear swelling in fl/fl and F-KO mice over 6 days (*n* = 4–6). (K) FACS of splenic Th17 cells in IMQ-treated fl/fl and F-KO mice on day 6. A representative histogram (*left*) and the number of Th17 cells (*right*) are shown. Values are mean ± s.e.m.. Representative data of two (A, F) or three (E) experiments, results of one experiment (B, I–K), and combined results of two (C, G, H) are shown. Statistical analysis was performed using unpaired t test (G, K), Mann-Whitney test (A, E, F, H), one-way ANOVA with Tukey’s multiple comparisons test (B, I), and two-way repeated measures ANOVA with Sidak’s multiple comparisons test (C, J). *, *P* < 0.05; **, *P* < 0.01.

To assess whether PLA2G12A expressed only in T cells or also in other cells would be important for Th17 differentiation, we performed a bone marrow (BM) chimera experiment in which irradiated *Pla2g12a*^+/+^ or *Pla2g12a*^-/-^ mice (recipients) were transplanted with *Pla2g12a*^+/+^ or *Pla2g12a*^-/-^ BM cells (donors) and then subjected to the psoriasis model (Figure S3A). Regardless of the recipient genotypes, IMQ-induced ear swelling (Figure S3B) and the proportions of Th17 and γδ T cells in the skin (Figures S3C and S3D) were markedly reduced when PLA2G12A was absent in the donor BM cells, consistent with the major contribution of PLA2G12A in T cells to the psoriasis model (Figures 3A–3G). Notably, transfer of *Pla2g12a*^+/+^ BM cells into *Pla2g12a*^-/-^ mice also partially reduced ear swelling (Figure S3B) and the proportions of Th17 and γδ T cells in the skin (Figures S3C and S3D), suggesting that PLA2G12A expressed in radioresistant non-hematopoietic cells was also partially involved in this event.

We further addressed the potential role of PLA2G12A expressed in non-T cells. *Pla2g12a* was expressed more abundantly in CD140a^+^ fibroblasts than in CD45^+^ hematopoietic cells in the LNs (Figure 3H). We therefore crossed *Pla2g12a*^fl/fl^ mice with transgenic mice overexpressing Cre recombinase under the *Col1a2* promoter to obtain fibroblast-specific PLA2G12A-deficient (*Pla2g12a*^fl/fl^*Col1a2*^cre^) mice. Expression of *Pla2g12a* was reduced by ∼70% in the whole LN tissue and by nearly 100% in CD140a^+^ fibroblasts, but not in CD45^+^ hematopoietic cells, within the LNs of *Pla2g12a*^fl/fl^*Col1a2*^cre^ mice relative to control mice (Figure 3H), confirming successful ablation of PLA2G12A in fibroblasts. *Ex vivo* Th17 differentiation of naïve T cells was reduced by nearly half in *Pla2g12a*^fl/fl^*Col1a2*^cre^ mice relative to *Pla2g12a*^fl/fl^ mice (Figure 3I). Accordingly, after IMQ treatment, *Pla2g12a*^fl/fl^*Col1a2*^cre^ mice showed significant reductions of ear swelling (Figure 3J) and splenic Th17 cells (Figure 3K) relative to *Pla2g12a*^fl/fl^ mice. Collectively, these results suggest that PLA2G12A secreted from Th17 cells critically contributes to psoriasis-like pathology and that PLA2G12A secreted from fibroblasts also has a supporting role possibly through its paracrine action on T cells.

### PLA2G12A deficiency attenuates arthritis

We next investigated the effect of PLA2G12A deficiency on collagen-induced arthritis (CIA), a well-established model of human rheumatoid arthritis that depends on Th17 immunity ^37^. For this purpose, we backcrossed *Pla2g12a*^-/-^ C57BL/6 mice onto a DBA/1 background, a strain that is sensitive to this model ^38^. The experimental procedure is depicted in Figure 4A. After the second immunization with type II collagen on day 21, *Pla2g12a*^+/+^ mice showed severe hindlimb swelling and an increased clinical score, whereas these symptoms were much milder in *Pla2g12a*^-/-^ mice (Figures 4B–4D). Histological analysis of the knee joints showed lower bone erosion and inflammatory cell infiltration, with weaker tartrate-resistant acid phosphatase (TRAP) staining suggestive of reduced osteoclast differentiation and activity, in *Pla2g12a*^-/-^ mice than in *Pla2g12a*^+/+^ mice (Figure 4E). Microcomputed tomography (μCT) analysis revealed that bone erosion, trabecular bone loss, and reduced cortical bone mineral density were less severe in *Pla2g12a*^-/-^ mice than in *Pla2g12a*^+/+^ mice (Figures 4F and 4G). Moreover, the increases in splenic Th17 cells (Figure 4H) and the serum IL-17A level (Figure 4I) following CIA were markedly attenuated by PLA2G12A deficiency. However, serum titers of anti-type II collagen antibody did not differ between the genotypes (Figure 4J), suggesting that loss of PLA2G12A did not profoundly affect humoral immune responses such as antigen uptake and presentation by DCs and antibody production by B cells.

**Figure 4.**
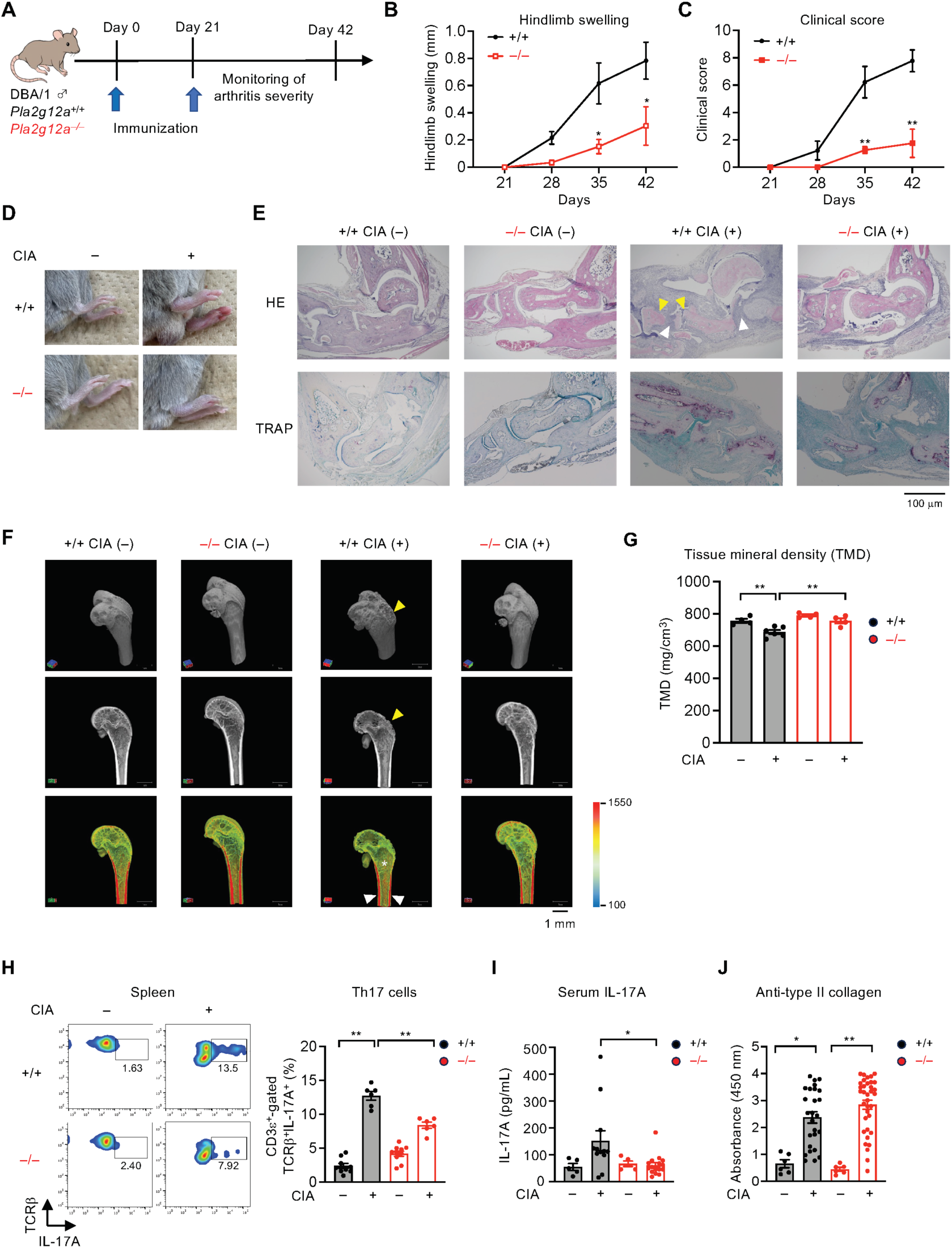
Amelioration of CIA by PLA2G12A deficiency. (A) Schematic representation of the procedure for IMQ-induced psoriasis. (B, C) Hindlimb swelling (*n* = 8–18) (B) and clinical score (n = 4–9) (C) in *Pla2g12a*^+/+^ (+/+) and *Pla2g12a*^-/-^ (–/–) mice over 42 days after immunization with type II collagen. (D) Representative photos of the hindlimbs of *Pla2g12a*^+/+^ and *Pla2g12a*^-/-^ mice with or without CIA on day 42. (E) HE and TRAP staining of the knee joints from *Pla2g12a*^+/+^ and *Pla2g12a*^-/-^ mice with (+) or without (–) CIA on day 42. Yellow arrowheads, bone erosion; white arrowheads, inflammatory cell infiltration. Scale bar, 100 µm. (F, G) μCT analysis of the hindlimb bone from *Pla2g12a*^+/+^ and *Pla2g12a*^-/-^ mice with or without CIA on day 42. Yellow arrowheads, bone erosion; white arrowheads, cortical bone; asterisk, trabecular bone. Scale bar, 1 mm. Representative μCT images (F) and tissue mineral density of the cortical bone (G) are shown. (H) FACS of splenic Th17 cells in *Pla2g12a*^+/+^ and *Pla2g12a*^-/-^ mice with or without CIA on day 42. Representative FACS profiles (*left*) and the proportion of Th17 cells (*right*) are shown. (I, J) ELISA of IL-17A (I) and anti-type II collagen (J) in sera of *Pla2g12a*^+/+^ and *Pla2g12a*^-/-^ mice with or without CIA on day 42. Values are mean ± s.e.m.. Representative data of three (B, C, G) or two (H) experiments and combined results of two experiments (I, J) are shown. Statistical analysis was performed using one-way ANOVA with Tukey’s multiple comparisons test (G, H), Kruskal-Wallis with Dunn’s multiple comparisons test (I, J), and two-way repeated measures ANOVA with Sidak’s multiple comparisons test (B, C). *, *P* < 0.05; **, *P* < 0.01.

To further confirm the contribution of PLA2G12A to arthritis on the C57BL/6 background, we employed a K/BxN serum transfer model of inflammatory arthritis, which also depends on IL-17A ^39^ and can be induced in various mouse strains ^40^. Transfer of the K/BxN mouse serum (using the procedure shown in Figure S4A) increased ankle thickness and clinical score peaking on day 4 and then declining to basal levels by day 11, being significantly milder in *Pla2g12a*^-/-^ mice than in *Pla2g12a*^+/+^ mice (Figures S4B and S4C). qPCR of the knee joints showed that the K/BxN-induced expression of *Il17a* was reduced in *Pla2g12a*^-/-^ mice, whereas that of *Il6* and *Il23*, which were likely expressed in DCs, macrophages, or stromal cells, was unaffected (Figure S4D). Expression of the arthritis markers *Tnf* and *Ptgs2* was decreased in *Pla2g12a*^-/-^ mice relative to *Pla2g12a*^+/+^ mice (Figure S4D), verifying that absence of PLA2G12A ameliorates arthritis severity. Overall, these data provide an additional line of evidence that PLA2G12A promotes autoimmune arthritis by inducing IL-17A-producing T cells.

### The PLA2G12A–LPA axis is crucial for Th17 differentiation

The above results suggest that some lipid products mobilized by PLA2G12A might contribute to the regulation of Th17 cells. Since the enzymatic activity of PLA2G12A is controversial ^31,32^, we reevaluated it using a natural membrane assay, in which a given sPLA_2_ is incubated with tissue-extracted phospholipids as substrates to monitor the release of fatty acids and lysophospholipids by liquid chromatography coupled with electrospray inonization**–**tandem mass spectrometry (LC-ESI-MS/MS) ^41,42^. Incubation of phospholipids extracted from mouse skin with recombinant human PLA2G12A resulted in dose-dependent increases of various LPC and LPE species with an *sn*-1 SFA or MUFA (Figure S5A) and free MUFA/PUFAs including OA, linoleic acid (LA; 18:2), AA, eicosapentaenoic acid (EPA; ω3 22:5), and docosahexaenoic acid (DHA; ω3 22:6) (Figure S5B). However, PLA2G12A barely increased LPA, lysophosphatidylserine (LPS), lysophosphatidylglycerol (LPG), and lysophosphatidylinositol (LPI) in this assay (Figure S5A). Thus, PLA2G12A has the enzymatic capacity to preferentially hydrolyze phosphatidylcholine (PC) and phosphatidylethanolamine (PE) to generate LPC and LPE as well as free MUFA/PUFAs *in vitro*.

Having confirmed the enzymatic properties of PLA2G12A, we performed targeted lipidomics of lysophospholipids (with 16:0, 18:0 or 18:1) and PUFA metabolites, which are typical PLA_2_ products, in the LNs of *Pla2g12a*^+/+^ and *Pla2g12a*^-/-^ mice with or without IMQ challenge on day 1 (Figures 5A, 5B, and S6A), at which time *Pla2g12a* expression reached its peak (Figure 2B). Heatmap (Figure 5A) and quantified values (Figure 5B) of lysophospholipids in the LNs showed that two LPA species, LPA18:0 and LPA18:1, were highly elevated in *Pla2g12a*^+/+^ mice after IMQ challenge, whereas this event was not evident in *Pla2g12a*^-/-^ mice. Additionally, the LN levels of LPE species, which were nearly constant regardless of IMQ treatment, were substantially lower in *Pla2g12a*^-/-^ mice than in *Pla2g12a*^+/+^ mice (Figures 5A and 5B). LPA16:0 and LPC species also showed a similar trend (Figures 5A and 5B). In contrast, the LN levels of LPG, LPI, and LPS (Figures 5A and 5B), as well as those of fatty acid metabolites (Figures S6A), were not significantly altered by PLA2G12A deficiency. In the skin, none of the lysophospholipids differed significantly between the genotypes, with a trend toward slight decreases in several lysophospholipids such as LPC, LPE and LPA with 18:1 in *Pla2g12a*^-/-^ mice (Figure S6B). These results suggest that PLA2G12A mobilizes LPC, LPE and LPA mainly in the LNs during IMQ-induced psoriasis.

**Figure 5.**
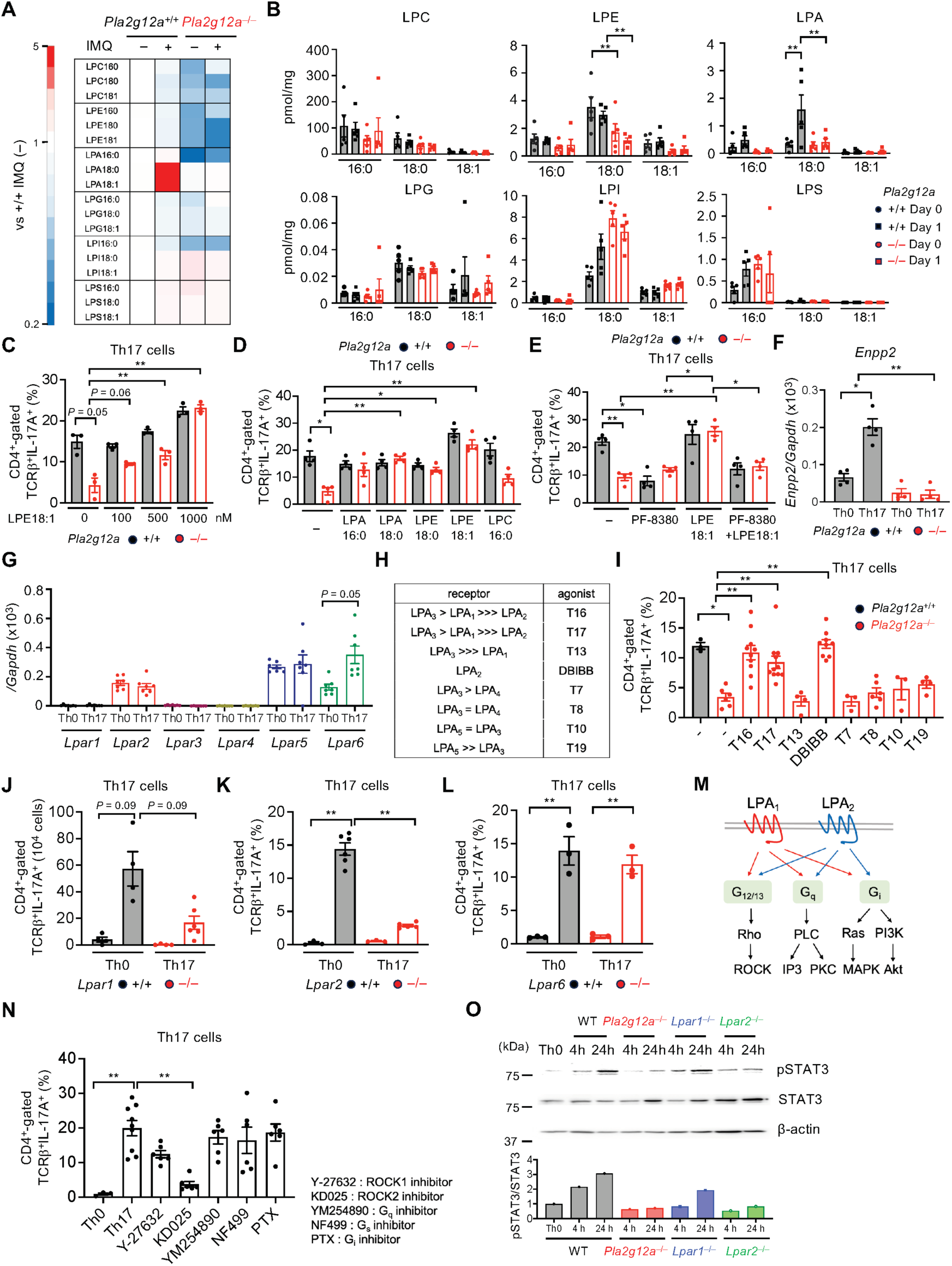
Mobilization of LPA by PLA2G12A is crucial for Th17 differentiation. (A, B) Lipidomics analysis of various lysophospholipids in the LNs of *Pla2g12a*^+/+^ (+/+) and *Pla2g12a*^-/-^ (–/–) mice with (+) or without (–) IMQ treatment for 1 day. Heatmap of individual lysophospholipids, with their levels in *Pla2g12a*^+/+^ mice without IMQ treatment as 1 (A), and their quantified values (pmol per mg tissue) (B) are shown. (C, D) FACS of Th17 cells after culture of *Pla2g12a*^+/+^ and *Pla2g12a*^-/-^ naïve T cells for 3 days with various concentrations of LPE(1-18:1) (C) or various lysophospholipids at 1 μM (D). (E) FACS of Th17 cells after culture of *Pla2g12a*^+/+^ and *Pla2g12a*^-/-^ naïve T cells for 3 days with the ATX inhibitor PF-8380 at 1 nM in the presence or absence of LPE(1-18:1). (F) qPCR of *Enpp2* in Th0 and Th17 cells of *Pla2g12a*^+/+^ and *Pla2g12a*^-/-^ mice on day 3. (G) qPCR of LPA receptors in Th0 and Th17 cells after culture for 3 days. (H) A list of LPA receptor agonists used in (I). (I) FACS of Th17 cells after culture of *Pla2g12a*^+/+^ and *Pla2g12a*^-/-^ naïve T cells for 3 days with various LPA receptor agonists at 10–100 nM. (J–L) FACS of Th17 cells after culture of naïve T cells prepared from *Lpar1*^+/+^ and *Lpar1^-/-^* mice (J), *Lpar2*^+/+^ and *Lpar2*^-/-^ mice (K), and *Lpar6*^+/+^ and *Lpar6*^-/-^ mice (L) for 3 days under Th0 and Th17 differentiation conditions. (M) A schematic illustration of LPA_1_ and LPA_2_ signaling. Both LPA_1_ and LPA_2_ can be coupled with G_12/13_, G_q_, and G_i_ signaling. (N) FACS analysis of Th17 cells after culture of WT naïve T cells for 3 days in the presence of inhibitors for G_12/13_, G_q_, G_i_ and G_s_ signaling. (O) Immunoblotting of STAT3, p-STAT3, and β-actin. Representative immunoblots (*upper*) and quantification of the p-STAT3/STAT3 ratio (*lower*) are shown. Values are mean ± s.e.m.. A result of one experiment (L), representative data of two experiments (A–G, J, K, N, O), and combined results of two experiments (I) are shown. Statistical analysis was performed using one-way ANOVA with Tukey’s multiple comparisons test (E, L), Brown-Forsythe and Welch ANOVA with Dunnett’s T3 multiple comparisons test (C, D, F, G, I–K, L), and two-way ANOVA with Sidak’s multiple comparisons test (B). *, *P* < 0.05; **, *P* < 0.01.

In agreement with our previous study ^29^, addition of LPE(1-18:1), an RORγt activator, to *Pla2g12a*^-/-^ T cells *ex vivo* restored Th17 differentiation in a dose-dependent manner (Figure 5C). In addition, the defective Th17 differentiation of *Pla2g12a*^-/-^ T cells was almost fully rescued by supplementation with LPA16:0, LPA18:0 or LPE18:0 and partially with LPC16:0 (Figure 5D). Since PLA2G12A hardly produced LPA *in vitro* (Figure S5A), we speculated that the LPE and LPC produced by PLA2G12A would be converted to LPA by autotaxin (ATX), a secreted lysophospholipase D that is abundantly present in culture medium containing serum ^43^, during Th17 culture. Indeed, the ATX inhibitor PF-8380 prevented Th17 differentiation of *Pla2g12a*^+/+^ cells and abrogated the restoring effect of LPE(1-18:1) on Th17 differentiation of *Pla2g12a*^-/-^ cells (Figure 5E). Moreover, expression of *Enpp2* (encoding ATX) was induced in *Pla2g12a*^+/+^, but not *Pla2g12a*^-/-^, Th17 cells over Th0 cells (Figure 5F). Thus, the conversion of LPE and likely LPC to LPA by ATX is crucial for Th17 differentiation.

LPA exerts its pluripotent biological effects by acting on six LPA receptors (LPA_1-6;_ encoded by *Lpar1-6*) ^44-46^, among which *Lpar2*, *Lpar5* and *Lpar6* were expressed at much higher levels than were *Lpar1, Lpar3* and *Lpar4* in both Th0 and Th17 cells (Figure 5G). This expression profile of LPA receptors in T cells is consistent with previous reports ^47,48^. To investigate the roles of LPA receptors in Th17 differentiation, we used a series of LPA receptor agonists ^49,50^ in Th17 differentiation culture (Figure 5H). Among them, the defective Th17 differentiation of *Pla2g12a*^-/-^ cells was restored by an LPA_2_-selective agonist (DBIBB) and LPA_1/3_-selective agonists (T16 and T17) (Figure 5I). Although T cells showed substantial expression of *Lpar5* (Figure 5H), which reportedly regulates CD8^+^ T cells in the context of anti-tumor immunity ^47,51,52^, LPA_5_-selective agonists (T10 and T19) were unable to rescue the defective Th17 differentiation of *Pla2g12a*^-/-^ cells (Figure 5I). Moreover, splenic naïve T cells prepared from LPA_1_-deficient (*Lpar1*^-/-^) or LPA_2_-deficient (*Lpar2*^-/-^) mice failed to differentiate properly into Th17 cells *ex vivo* (Figures 5J and 5K). Although expression of *Lpar6* was increased in Th17 cells relative to Th0 cells (Figure 5G), Th17 differentiation of native T cells from *Lpar6*^-/-^ mice was comparable to that of such cells from control mice (Figure 5L). Overall, in addition to LPE(1-18:1), which activates RORγt ^29^, LPA, which is produced by the sequential action of PLA2G12A and ATX, may be required for proper Th17 differentiation via LPA_1_ and/or LPA_2_ receptors.

Both LPA_1_ and LPA_2_ are coupled with G_i_, G_q_ and G_12/13_ signaling (Figure 5M) ^53^. Th17 differentiation of naïve T cells from WT mice was suppressed markedly by KD025 (ROCK2 inhibitor) and modestly by Y-27632 (ROCK1 inhibitor), which can dampen G_12/13_ signaling ^54,55^, but not by G_i_, G_q_ and G_s_ inhibitors (Figure 5N), suggesting that LPA_1_ and/or LPA_2_ are coupled mainly with the G_12/13_-Rho-ROCK2 signaling pathway in the context of Th17 differentiation. Phosphorylation of STAT3, a transcription factor that is essential for IL-6 and IL-23 signaling ^56^ and interacts with ROCK2 in Th17 cells ^57^, was markedly reduced in T cells from *Pla2g12a*^-/-^ or *Lpar2*^-/-^ mice, but only partially in those from *Lpar1*^-/-^ mice, in Th17 differentiation culture (Figures 5O). Collectively, downstream of PLA2G12A and ATX, the LPA_2_-G_12/13_-ROCK2 signaling pathway may be mainly responsible for the regulation of Th17 differentiation by enhancing STAT3 phosphorylation, while LPA_1_ may play a minor role in this context or be involved in some other regulatory steps.

While LPA_1_ deficiency hampered Th17 differentiation *ex vivo* (Figure 5J) and LPA_1/3_ agonists rescued the Th17 differentiation of *Pla2g12a*^-/-^ T cells (Figure 5I), addition of the LPA_1/3_ antagonist Ki16425 to the *ex vivo* culture revealed only a weak tendency to suppress Th17 differentiation (Figure S7A). Since the expression of *Lpar1* in Th17 cells at 72 h of culture was nearly negligible (Figure 5G), we examined the time course of its expression throughout the Th17 differentiation culture and found that it was substantially detectable at the earliest time point (0-6 h) of both Th0 and Th17 cultures, rapidly declining thereafter (Figure S7B). Thus, LPA_1_ appears to be downregulated after TCR activation and to work only during the initial phase of Th17 differentiation. This may explain why the strong activation of LPA_1_ expressed during the early period of culture by its agonists was able to overcome the defective Th17 differentiation of *Pla2g12a*^-/-^ cells, whereas this was poorly suppressed by an LPA_1/3_ antagonist in *Pla2g12a*^+/+^ cells in the continued presence of sufficient LPA_2_ signal.

Global deficiency of LPA_1_ exacerbated, rather than ameliorated, IMQ-induced ear swelling (Figures S7C–S7E), possibly because LPA_1_ expressed in dermal fibroblasts, epidermal keratinocytes, or other cells has skin-intrinsic protective effects ^42,58,59^. However, adoptive transfer of BM cells from *Lpar1*^-/-^ mice into irradiated WT mice resulted in reductions of IMQ-induced ear swelling (Figure S7F) and splenic Th17 induction (Figure S7G), relative to that elicited by BM cells from *Lpar1*^+/+^ mice. Notably, under this condition, the proportion of splenic CD3ε^+^ T cells was lower in mice receiving *Lpar1*^-/-^ BM cells than in those receiving *Lpar1*^+/+^ BM cells (Figure S7H). In contrast, in mice receiving *Pla2g12a*^-/-^ BM cells, the proportion of splenic CD3ε^+^ T cells was unaffected (Figure S7I). Moreover, under Th0 culture conditions, LPA_1_ deficiency reduced the proportion of CD4^+^ T cells, while neither PLA2G12A nor LPA_2_ deficiency profoundly altered it (Figures S7J), suggesting that LPA_1_ regulates surface CD4 expression independently of PLA2G12A and LPA_2_. Taken together, these findings suggest that LPA_2_ likely acts as a main LPA receptor that assists Th17 differentiation downstream of PLA2G12A, whereas LPA_1_ is more likely to be involved in the regulation of T cell development or activation, which secondarily affects Th17 differentiation. In addition, among the splenic CD4^+^ T cell population, the ratio of CD44^+^CD62L^-^ effector memory CD4^+^ T cells to CD44^-^CD62L^+^ naïve CD4^+^ T cells was reduced to a similar degree in *Pla2g12a*^-/-^ and *Lpar2*^-/-^ mice, but only modestly in *Lpar1*^-/-^ mice (Figure S7K), further supporting the functional association of PLA2G12A with LPA_2_ rather than with LPA_1_.

### PLA2G12A modifies Th17-derived EVs

A growing body of evidence suggests that sPLA_2_s hydrolyze phospholipids in EV membranes to release bioactive lipids ^42,60,61^. Small EVs, often referred to as exosomes, are produced by and/or act on T cells, thereby variably influencing T cell functions ^62,63^. We noted that *ex vivo* differentiated Th17 cells, but not Th0 cells, from *Pla2g12a*^+/+^ mice secreted small EVs of 50∼150 nm in diameter (relevant to the size of exosomes) (Figures S8A-S8E), as reported previously ^64^. The secretion of EVs (bulk small EVs and CD9^+^CD63^+^ EVs) from Th17 cells was markedly reduced in *Pla2g12a*^-/-^ mice as well as in *Lpar1*^-/-^ and *Lpar2*^-/-^ mice relative to WT mice (Figures S8A and S8B). Transmission electron microscopy (TEM) and cryogenic electron microscopy (cryo-EM) revealed that although the morphology of EVs prepared from *Pla2g12a*^-/-^ Th17 cells (*Pla2g12a*^-/-^ EVs) was similar to that of EVs from *Pla2g12a*^+/+^ Th17 cells (*Pla2g12a*^+/+^ EVs), *Pla2g12a*^-/-^ EVs tended to have a broader size distribution than did *Pla2g12a*^+/+^ EVs (Figures S8C-S8E). Moreover, treatment of these EVs with recombinant PLA2G12A reduced their average particle sizes (Figures S8C-S8E), in accord with a previous observation that lymphoma-derived EVs became smaller after treatment with PLA2G10 (sPLA_2_-X) ^61^. In line with the reduced secretion of EVs by *Pla2g12a*^-/-^ Th17 cells, microarray analysis revealed that the expression levels of a set of genes related to EV formation and secretion were increased in *Pla2g12a*^+/+^, but not *Pla2g12a*^-/-^, Th17 cells relative to Th0 cells (Figure S8F). Thus, PLA2G12A affects the secretion and size of Th17-derived EVs.

Lipidomics analysis of *Pla2g12a*^+/+^ and *Pla2g12a*^-/-^ EVs revealed that their overall phospholipid composition, except for PE, was similar (Figure S8G). For PE, there were apparent increases in the proportions of PE34:1 and PE36:1, which have an OA (18:1) chain, in *Pla2g12a*^-/-^ EVs relative to *Pla2g12a*^+/+^ EVs (Figure S8G), likely mirroring the reduced hydrolysis of these PE species by endogenous PLA2G12A. Importantly, the amounts of LPE and LPC species, including LPE(1-18:1), were markedly lower in *Pla2g12a*^-/-^ EVs than in *Pla2g12a*^+/+^ EVs (Figure 6A), implying that PLA2G12A acts on Th17-derived EVs to generate LPE and LPC. However, LPA levels in *Pla2g12a*^+/+^ and *Pla2g12a*^-/-^ EVs were similar (Figure 6A), suggesting that the conversion of LPE and LPC to LPA by ATX may take place outside the EVs or that ATX may be supplied mainly from other sources, such as stromal fibroblasts ^42^ or lymphatic endothelial cells ^65-67^, under *in vivo* conditions. Addition of recombinant PLA2G12A to *Pla2g12a*^+/+^ EVs increased the levels of LPE and LPC (particularly the former) (Figure 6B), further supporting the notion that PLA2G12A targets EV membranes.

**Figure 6.**
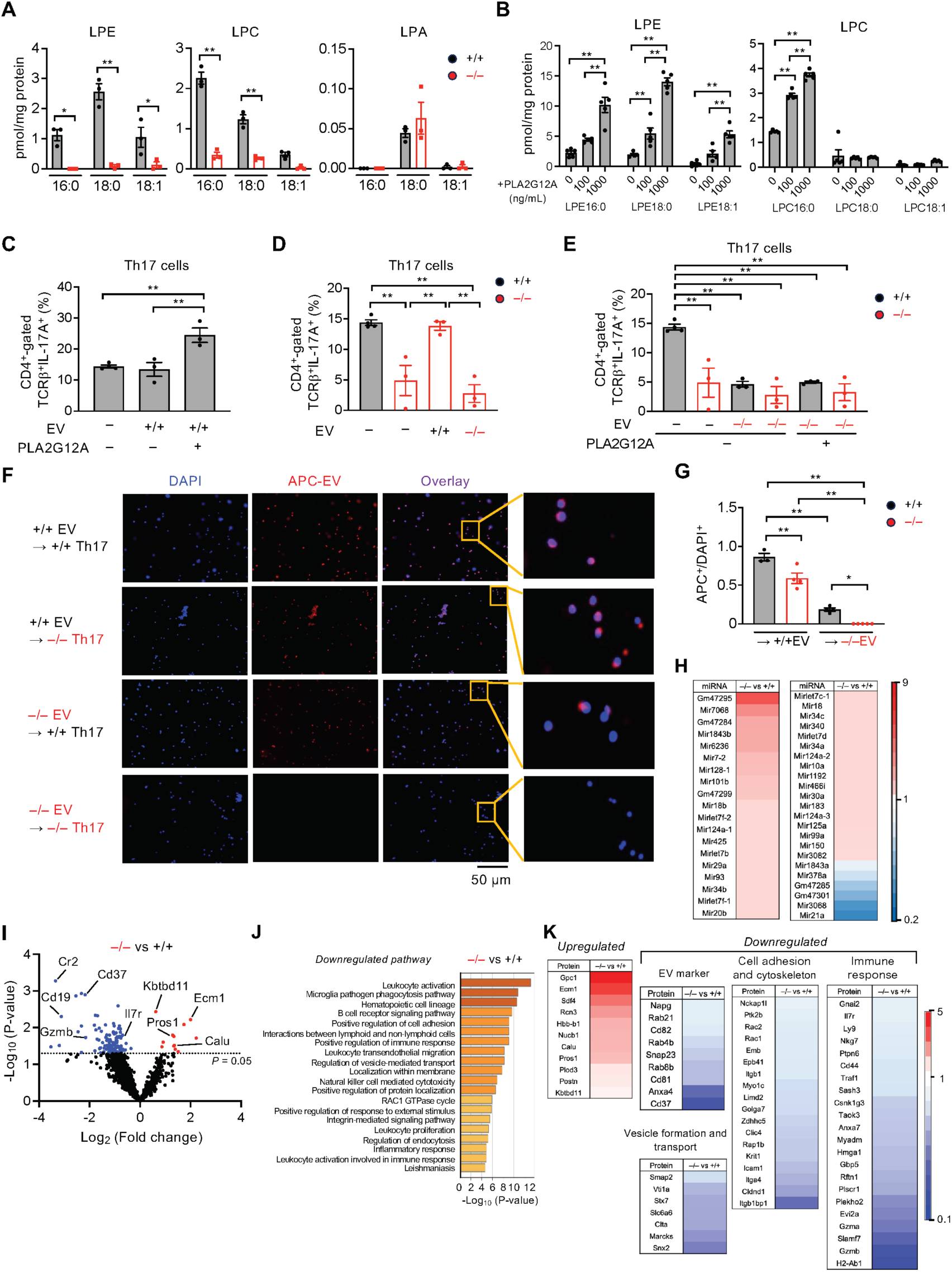
Alterations in the properties and functions of Th17-derived EVs by PLA2G12A deficiency. (A) Lysophospholipid levels in EVs isolated from *Pla2g12a*^+/+^ (+/+) and *Pla2g12a*^-/-^ (–/–) Th17 cells after culture for 3 days. (B) LPE and LPC levels after incubation of *Pla2g12a*^+/+^ Th17-derived EVs (+/+ EVs) for 4 h with the indicated concentrations of recombinant mouse PLA2G12A. (C) FACS of Th17 cells after culture of *Pla2g12a*^+/+^ naïve T cells for 3 days in the presence or absence of +/+ EVs that had been treated with or without 1 μg/ml PLA2G12A. (D) FACS of Th17 cells after culture of naïve T cells from *Pla2g12a*^+/+^ and *Pla2g12a*^-/-^ mice for 3 days in the presence or absence of +/+ EVs or *Pla2g12a*^-/-^ Th17-derived EVs (–/– EVs). (E) FACS of Th17 cells after culture of naïve T cells from *Pla2g12a*^+/+^ and *Pla2g12a*^-/-^ mice for 3 days in the presence or absence of –/– EVs that had been treated with or without the indicated concentrations of PLA2G12A. (F, G) Immunofluorescence microscopy of WT and *Pla2g12a*^-/-^ Th17 cells cultured for 4 h with APC-labelled +/+ EVs or –/– EVs. Representative images (F) and the ratio of EV-captured (APC^+^) cells to all (DAPI^+^) cells (G) are shown. Scale bar, 50 µm. (H) Heatmap of representative miRNAs whose levels were altered in –/– EVs relative to +/+ EVs. Colors indicate fold changes in –/– EVs relative to +/+ EVs. (I–K) Proteome analysis of EVs. Volcano plot (I), pathway enrichment analysis (J), and heatmap (K) of representative proteins whose levels were altered in +/+ EVs relative to –/– EVs. Colors in (K) indicate fold changes in –/– EVs relative to +/+ EVs. Values are mean ± s.e.m.. Representative data of two experiments (A) and results of one experiment (B–K) are shown. Statistical analysis was performed using two-way ANOVA with Sidak’s multiple comparisons test (A, B) and ordinary one-way ANOVA with Tukey’s multiple comparisons test (C-E, G). *, *P* < 0.05; **, *P* < 0.01.

Addition of PLA2G12A-treated EVs, with enriched LPE(1-18:1), to *Pla2g12a*^+/+^ naïve T cells in the Th17 differentiation culture augmented Th17 differentiation (Figure 6C). Defective Th17 differentiation of *Pla2g12a*^-/-^ cells was almost fully rescued by supplementation with *Pla2g12a*^+/+^ EVs, but not with *Pla2g12a*^-/-^ EVs (Figure 6D). Moreover, *Pla2g12a*^-/-^ EVs hindered the Th17 differentiation of *Pla2g12a*^+/+^ T cells to a level similar to that of *Pla2g12a*^-/-^ T cells, even after pretreatment with PLA2G12A (Figure 6E). These results suggest that, beyond phospholipid hydrolysis, *Pla2g12a*^+/+^ EVs and *Pla2g12a*^-/-^ EVs have immunostimulatory and immunosuppressive features, respectively, in the context of Th17 differentiation. PLA2G10-modified lymphoma EVs are readily incorporated and deliver their cargos to target macrophages ^61^. In line with this, immunofluorescence microscopy showed that allophycocyanin (APC)-labeled *Pla2g12a*^+/+^ EVs were taken up into the cytoplasm of *Pla2g12a*^+/+^ Th17 cells and to a slightly lesser extent that of *Pla2g12a*^-/-^ Th17 cells, whereas uptake of *Pla2g12a*^-/-^ EVs was strikingly reduced in *Pla2g12a*^+/+^ Th17 cells and reduced even further in *Pla2g12a*^-/-^ Th17 cells (Figures 6F and 6G). Thus, the uptake of EVs and therefore the transfer of EV cargos to Th17 cells are markedly impaired by PLA2G12A deficiency.

To further characterize the alterations in EV cargos, we performed RNA sequencing (RNA-seq) and proteome analyses of *Pla2g12a*^+/+^ and *Pla2g12a*^-/-^ EVs. Although the overall RNA cargo profiles were not changed dramatically by PLA2G12A deficiency, the abundance of specific microRNAs (miRNAs) and other RNA classes apparently differed between *Pla2g12a*^+/+^ and *Pla2g12a*^-/-^ EVs (Figures 7H and S9A-S9C). Focusing on miRNAs that have been implicated in the regulation of Th17 immunity ^68^, the level of miR-21, which enhances Th17 differentiation by targeting Smad7 ^69^, was markedly lower, whereas the levels of miR-20, -29, -30, -34, -93,-124, and -3082, which attenuate Th17 signaling by targeting Th17-related cytokines, their receptors, and transcription factors ^70-75^, and miR-let7s, which repress effector T cell functions ^76^, were higher, in *Pla2g12a*^-/-^ EVs than in *Pla2g12a*^+/+^ EVs (Figure 6H). The changes in these miRNA cargos could account, at least in part, for the Th17-suppressing effect of *Pla2g12a*^-/-^ EVs. A volcano plot of the EV proteome also revealed a striking difference in the abundance of cargo proteins between *Pla2g12a*^+/+^ and *Pla2g12a*^-/-^ EVs (Figure 6I). Pathway enrichment analysis showed that proteins related to EV markers (*e.g.*, CD19, Anxa4, and Rab proteins), vesicle formation and transport (*e.g*., Marcks, Clta, and Snx2), cell adhesion and cytoskeleton (*e.g.*, claudin, ICAM, integrins, and Rac proteins), and immune responses (*e.g*., Gnai2, IL-7R, and Rftn1) were reduced, whereas proteins that potentially modulate immune responses (*e.g*., Gpc1, Ecm1, Pros1, Rcn3, and Plod3) were increased, in *Pla2g12a*^-/-^ EVs relative to *Pla2g12a*^+/+^ EVs (Figures 6J and 6K), which could also be associated with the immunostimulatory and immunosuppressive aspects of *Pla2g12a*^+/+^ and *Pla2g12a*^-/-^ EVs, respectively. Overall, PLA2G12A alters EV cargos including lipids, RNAs and proteins that could influence the secretion, uptake, and functions of EVs, thereby controlling Th17 differentiation.

**Figure 7.**
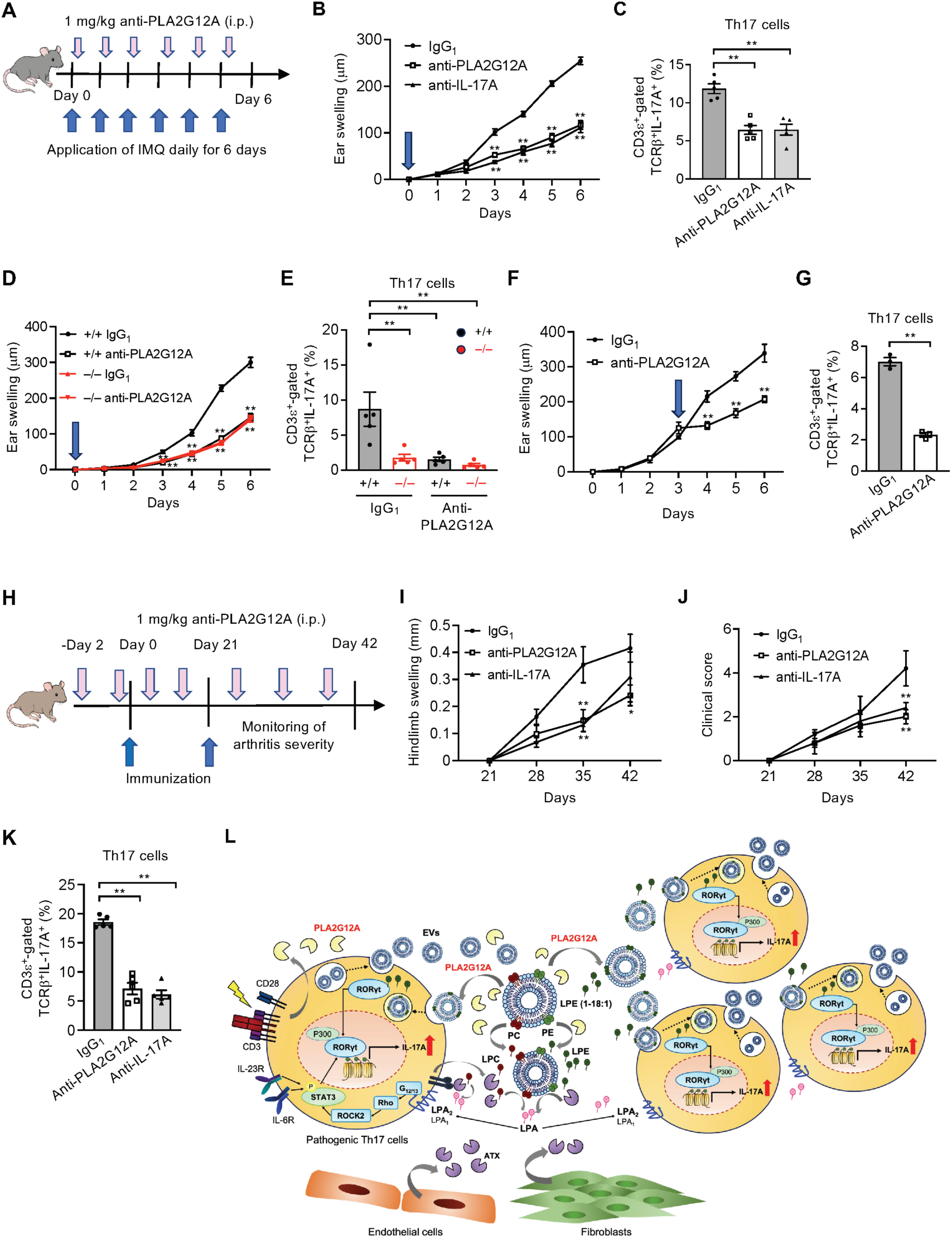
Anti-PLA2G12A antibody ameliorates psoriasis and arthritis models. (A) Schematic representation of the procedure used for antibody treatment in IMQ-induced psoriasis. (B) IMQ-induced ear swelling in WT mice treated with anti-PLA2G12A, -IL-17A, or control IgG_1_ antibody over 6 days. Antibody treatment started on day 0 (arrow). (C) FACS of splenic Th17 cells in (B) on day 6. (D) IMQ-induced ear swelling in *Pla2g12a*^+/+^ (+/+) and *Pla2g12a*^-/-^ (–/–) mice treated with anti-PLA2G12A or control antibody over 6 days. Antibody treatment started on day 0 (arrow). (E) FACS of splenic Th17 cells in (D) on day 6. (F) IMQ-induced ear swelling in WT mice treated with anti-PLA2G12A or control antibody. Antibody treatment started on day 3 (arrow). (G) FACS of splenic Th17 cells in (F) on day 6. (H) Schematic representation of the procedure used for antibody treatment in CIA. (I, J) Paw thickness (I) and clinical sore (J) in WT mice treated with anti-PLA2G12A, - IL-17A, or control antibody over 42 days. (K) FACS of splenic Th17 cells in (I, J) on day 42. (L) Schematic diagram showing PLA2G12A-driven regulation of Th17 differentiation. PLA2G12A is induced in CD4^+^ T cells after TCR stimulation and is also supplied from stromal fibroblasts. EV secretion is facilitated by Th17 differentiation driven by Th17-inducing cytokines. PLA2G12A hydrolyzes PE and PC to generate LPE and LPC in EV membranes. The EVs are taken up by Th17-differentiating cells and deliver LPE(1-18:1) intracellularly, which then transactivates RORγt. LPE and LPC are converted to LPA by ATX, which is supplied from Th17 cells, stromal cells (*e.g*., fibroblasts and endothelial cells), and serum. LPA acts mainly on LPA_2_ (and to a lesser extent LPA_1_) on Th17-differentiating cells to augment IL-6/IL23-mediated STAT3 phosphorylation via G_12/13_-Rho-ROCK2 signaling, assisting pathogenic Th17 differentiation in autocrine and paracrine fashions. Gene disruption of PLA2G12A dampens this lipid-orchestrated amplification loop and decreases EV secretion, uptake, and transfer of lipid, miRNA, and protein cargos. Accordingly, antibody-mediated neutralization of PLA2G12A efficiently attenuates psoriasis, arthritis, and probably other Th17-related pathologies such as EAE. Values are mean ± s.e.m.. Results are from one experiment (B–G, I–K). Statistical analysis was performed using unpaired t test (G), ordinary one-way ANOVA with Dunnett’s multiple comparisons test (C, E, K), and two-way repeated measures ANOVA with Sidak’s multiple comparisons test (B, D, F, I, J). *, *P* < 0.05; **, *P* < 0.01 *versus* control IgG_1_.

### Anti-PLA2G12A antibody attenuates psoriasis and arthritis

The results described above raised the possibility that pharmacological inhibition of PLA2G12A could attenuate Th17-related diseases. Since no agent that specifically inhibits PLA2G12A has currently been commercially available, we attempted to generate monoclonal antibodies capable of neutralizing this atypical sPLA_2_. To this end, we immunized *Pla2g12a*^-/-^ mice with recombinant human PLA2G12A-Fc fusion protein. After two rounds of screening, we selected six hybridoma clones producing IgG_1_ antibody that inhibited the enzymatic activity of both human and mouse PLA2G12A proteins in an *in vitro* enzyme assay (Figure S10A). These antibodies efficiently blocked the *ex vivo* differentiation of naïve T cells into Th17 cells (Figure S10B). We used the antibody clone #44, which showed the highest neutralizing activity, in subsequent studies. This antibody reacted with human and mouse PLA2G12A proteins, but not with other sPLA_2_ isoforms, on immunoblotting (Figure S10C) and ELISA (Figure S10D), confirming its specificity. Immunohistochemistry of mouse skin using the anti-PLA2G12A antibody revealed that dermal clusters of CD4^+^ T cells, as well as epidermal keratinocytes, dermal fibroblasts, and subcutaneous muscle layers, were stained in *Pla2g12a*^+/+^, but not *Pla2g12a*^-/-^, mice (Figure S10E). The CD4^+^ T cell clusters were focally increased in the dermis following IMQ treatment and were smaller in *Pla2g12a*^-/-^ mice than in *Pla2g12a*^+/+^ mice (Figure S10E), consistent with reduced psoriatic inflammation by PLA2G12A deficiency.

We then applied this anti-PLA2G12A antibody to the IMQ-induced psoriasis model. When 1 mg/kg anti-PLA2G12A antibody was intraperitoneally injected into C57BL/6 mice once daily for 6 days in accordance with the procedure shown in Figure 7A, ear swelling over time (Figure 7B) and the splenic proportion of Th17 cells on day 6 (Figure 7C) were reduced to levels similar to those in replicate mice treated with an anti-IL-17A antibody used as a positive control. The attenuation of psoriatic inflammation by the anti-PLA2G12A antibody was equivalent to that resulting from PLA2G12A deficiency (Figure 2). No further suppression was observed when *Pla2g12a*^-/-^ mice were treated with the anti-PLA2G12A antibody (Figures 7D and 7E), confirming that the therapeutic effect of the antibody was due to specific inhibition of PLA2G12A, rather than an off-target effect. In the same model, administration of the anti-PLA2G12A antibody into WT mice on day 3 halted subsequent disease progression (Figures 7F and 7G). We also applied the anti-PLA2G12A antibody to the CIA model. When 1 mg/kg anti-PLA2G12A antibody was injected intraperitoneally into DBA/1 mice once a week in accordance with the procedure shown in Figure 7H, paw thickness (Figure 7I), clinical score (Figure 7J), and splenic Th17 cells (Figure 7K) were reduced to levels comparable to those in replicate mice treated with an anti-IL-17A antibody. The suppressive effect of the anti-PLA2G12A antibody on arthritic inflammation was equivalent to that of PLA2G12A deficiency (Figures 4B, 4C and 4H). Thus, the anti-PLA2G12A antibody we generated in this study can efficiently block mouse models of psoriasis and arthritis.

## DISCUSSION

Accumulating evidence suggests that specific lipid-metabolic pathways for fatty acid- or cholesterol-related metabolites play an essential role in the differentiation and function of pathogenic Th17 cells ^10,12,17,23,26^. Through CRISPR-based knockout screening combined with comprehensive lipidomics of CD4^+^ T cells, we have recently reported that the *de novo* synthesis of LPE(1-18:1) through a series of specific lipogenic enzymes, including ACC1, FASN, SCD2 and GPAMs, is required for proper RORγt function directed toward pathogenic Th17 differentiation ^27,29^. Since the generation of lysophospholipids requires a lipolytic PLA_2_ reaction, it is likely that PLA2G12A, one of the lipid-metabolic enzymes identified by CRISPR screening thus far, is responsible for the generation of LPE(1-18:1) as an essential modulator of RORγt activity ^29^. Herein, we have extended this finding and provided more profound insights into the mechanistic action of this secreted enzyme during Th17 differentiation and Th17-associated pathology.

PLA2G12A, an atypical member of the sPLA_2_ family, is markedly induced in CD4^+^ T cells after TCR activation. Global, CD4^+^ T cell-specific, or fibroblast-specific ablation of PLA2G2A, as well as its antibody-based neutralization, attenuates the differentiation of IL-17A-producing T cells (Th17 and γδ T cells) and associated pathology. In addition to generating LPE(1-18:1) as an RORγt activator ^29^, PLA2G12A orchestrates a sophisticated extracellular lipid circuit via hydrolysis of EV membranes, which alters the properties and functions of the EVs and also supplies LPA, another bioactive lipid that acts as a supportive signal for proper Th17 differentiation. Thus, our present study has established the concept of the sPLA_2_-EV-lysophospholipid axis as a general mode of sPLA_2_ action, unveiled a T cell-intrinsic immunoregulatory role of the LPA_2_ receptor, and underpinned a critical role of EVs as carriers of unique lipids, miRNAs and proteins for the regulation of Th17 immunity. The overall scheme underlying the PLA2G12A-driven regulation of Th17 differentiation is summarized in Figure 7L.

In line with an emerging view that EV membranes serve as an unequivocal hydrolytic target for sPLA_2_s ^42,60,61,77,78^, our results suggest that PLA2G12A hydrolyzes PC and PE to give rise to LPC and LPE, including LPE(1-18:1), in Th17-derived EVs. T cell-derived EVs play various roles in cell-to-cell signaling in the context of inflammatory responses, autoimmunity, and infectious diseases ^62^. We show here that Th17 cells secrete EVs that can amplify their own differentiation and function and that this process depends on sPLA_2_-driven lipid signaling. The PLA2G12A-modified EVs are taken up by Th17 cells, allowing intracellular delivery of exogenously produced LPE(1-18:1), thus making it accessible to the nuclear receptor RORγt. Moreover, Th17-derived EVs contain miRNAs and proteins that can mediate EV binding to cells and activation of immune responses, likely conferring pathogenic potential on neighboring T cells for propagation of Th17 immunity. EVs secreted from PLA2G12A-deficient Th17 cells contain altered levels of specific miRNAs and proteins that potentially support or suppress Th17 differentiation, and accordingly, loss of PLA2G12A markedly diminishes the ability of EVs to deliver their cargos to recipient Th17 cells, eventually hampering proper Th17 differentiation. This is compatible with our previous finding that hydrolytic modification of EVs derived from B-cell lymphomas by PLA2G10 increases the intercellular transfer of viral miRNAs to recipient macrophages, allowing their polarization into an immunosuppressive and tumor-promoting phenotype ^61^. Importantly, promotion of Th17 differentiation by PLA2G12A ultimately increases the secretion of EVs as a hydrolytic platform for this sPLA_2_, constituting a feed-forward loop that further amplifies Th17 differentiation and related pathogenesis, and this may explain why PLA2G12A deficiency alters EV cargo contents in association with Th17 differentiation.

LPA acts as a pluripotent lysophospholipid mediator that regulates various biological processes including vascular development, hair follicle homeostasis, neuropathic pain, reproduction, and cancer ^44-46^. In T cell biology, LPA_5_ and LPA_6_ receptors suppress the infiltration or cytotoxic function of CD8^+^ T cells, thereby impeding anti-tumor immunity^47,48,51,52^. We have demonstrated here that PLA2G12A-driven Th17 differentiation requires LPA production. PLA2G12A secreted from Th17 cells acts on EV membranes to release LPE and LPC, which are then converted by ATX to LPA. Thus, the two secreted lipolytic enzymes, PLA2G12A and ATX, supplied from hematopietic and stromal compartments cooperatively produce LPA within a given extracellular milieu. Furthermore, our genetic and pharmacological approaches have revealed that LPA_2_, and to a lesser extent LPA_1_, are responsible for Th17 differentiation by augmenting STAT3 phosphorylation. Thus, EV hydrolysis by PLA2G12A mobilizes two particular bioactive lysophospholipids acting on Th17 cells: LPE(1-18:1), which is delivered intracellularly via EV uptake and activates the nuclear receptor RORγt, and LPA, which acts mainly on cell surface LPA_2_ to augment STAT3 activation. While the ATX-LPA_2_ axis reportedly regulates intranodal T cell motility via Gq-Rac signaling ^79,80^ and cell survival and invasion via G_12/13_-YAP signaling ^81-83^, our present study has revealed a role of LPA_2_- G_12/13_-ROCK2 signaling in the regulation of Th17 immunity. In line with our present findings, it has been reported that LPA_1_ deficiency alleviates a mouse arthritis model with reduced Th17 differentiation ^84^. However, the action of LPA_1_ is somehow complex in this context, since its expression in CD4^+^ T cells is low and precedes Th17 differentiation and since its loss seems to perturb T cell development or activation prior to Th17 differentiation. Moreover, LPA_1_ is expressed in various cell types ^42,58,59^ and its sytemic *versus* BM-specific deletions have different impacts on the psoriasis model. Although beyond the scope of the present study, clarification of the precise roles of LPA_1_ in T cell development and activation would be warranted.

Historically, PLA_2_ enzymes have long been implicated in the generation of fatty acid metabolites, especially AA-derived eicosanoids. However, our current studies have provided compelling evidence that sPLA_2_s are better suited for generation of bioactive lysophospholipids in extracelluler fluids. For instance, PLA2G3 (sPLA_2_-III)-driven LPA production in mast cell-derived EVs promotes mast cell maturation via fibroblast LPA_1_^42^, pulmonary PLA2G3-driven LPA generation protects against asthma via epithelial LPA_2_ ^85^, PLA2G10-driven LPA generation in lymphoma-derived EVs facililates macrophage polarization toward a pro-tumorigenic phenotype probably via LPA_1_ ^61^, and keratinocyte PLA2G2F (sPLA_2_-IIF)-driven generation of lysoplasmalogen (alkenyl-LPE) promotes epidermal hyperplasia via STAT3 activation ^86,87^. In this regard, PLA2G12A-mediated generation of LPE and LPA for Th17 differentiation represents another example of the sPLA_2_-EV-lysophospholipid axis for fine-tuning of tissue homeostasis and diseases, thus generalizing the action mode of the sPLA_2_ family.

Importantly, our study has provided a rationale for the possible use of PLA2G12A as a drug target for Th17-related disease. Current sPLA_2_ inhibitors, which block classical group I/II/V/X sPLA_2_s altogether because of the structural similarity of their catalytic pocket, have failed to improve the symptoms of rheumatoid arthritis, sepsis, asthma, and cardiovascular disease in clinical trials ^88-90^. This is most probably because blunting the functions of all classical sPLA_2_s by pan-sPLA_2_ inhibitors would cancel both the beneficial and detrimental effects of these enzymes and thereby yield no desirable pharmacological impact ^91^. Since none of the current sPLA_2_ inhibitors block PLA2G3 and PLA2G12A, whose structures differ markedly from those of classical group I/II/V/X sPLA_2_s, these atypical sPLA_2_s may be attractive drug targets with high specificity. Our present finding that a PLA2G12A-neutralizing monoclonal antibody was able to efficiently ameliorate psoriasis and arthritis models with an efficacy comparable to that of PLA2G12A deficiency or anti-IL-17A antibody treatment has opened a new avenue for development of new drugs targeting this unique sPLA_2_. Since T cell-specific PLA2G12A deletion also ameliorates experimental autoimmune encephalomyelitis (EAE), a model for human multiple sclerosis ^29^, the anti-PLA2G12A antibody could be useful for treatment of various Th17-related diseases. Although antibody-based medicines that target cytokines or their receptors have shown potent efficacy and are now used clinically to treat patients with psoriasis and rheumatoid arthritis ^92^, they have side-effects that increase the risk of infection and disruption of the epithelial barrier. Indeed, blocking IL-17A or its receptor effectively attenuates psoriasis, while worsening inflammatory bowel diseases due to disturbance of the gut mucosal barrier ^93^. Given that the PLA2G12A-driven lipid pathway facilitates pathogenic rather than homeostatic Th17 differentiation, the anti-PLA2G12A antibody may represent a new therapeutic agent for Th17-related diseases and even pave the way for future development of a small-molecule compound that would specifically inhibit this atypical sPLA_2_.

There are several remaining questions that need to be resolved in future studies. It remains unclear how PLA2G12A-hydrolyzed EVs are taken up by Th17 cells (possibly via clathrin- or caveolae-mediated endocytosis, macropinocytosis, membrane fusion, or other routes), how LPE(1-18:1) in the internalized EVs is presented to RORγt in the cytosol and/or nucleus, and which specific miRNAs or proteins in these EVs are truly responsible for the regulation of Th17 differentiation. Redundant or distinct roles of LPA_1_ and LPA_2_ in T cell development, activation, or polarization into Th17 cells, including the underlying signaling pathways therein, require further elucidation. The failure of LPA_6_, which is known to be coupled with G_12/13_, to participate in PLA2G12A-driven Th17 differentiation is probably because it recognizes LPA species with an *sn*-2 fatty acid (PLA_1_ product) in preference to those with an *sn*-1 fatty acid (PLA_2_ product) ^94^. Marked and transient upregulation of PLA2G12A following TCR activation suggests potential involvement of this sPLA_2_ in the regulation of T cell activation and even differentiation into other T cell subsets as well. Indeed, our preliminary study suggests that PLA2G12A deficiency also perturbs Th1 and Th2 differentiation, which should be clarified in the context of Th1/Th2-associated diseases in future studies. Although it is unclear how the lack of PLA2G12A decreases EV secretion from Th17 cells, this might be related to the observation that PLA2G12B, the closest homolog of PLA2G12A with no enzymatic activity ^95^, regulates lipoprotein secretion by acting as a lipid-assembling chaperone in hepatocytes ^96^. Considering the broad tissue distribution of PLA2G12A, the contributions of this sPLA_2_ to other biological events beyond T cell immunity, such as metabolic regulation, neuronal process, host defense, and cancer ^97-102^, should also be taken into consideration. Deciphering the enzymatic activity-dependent or -independent functions of PLA2G12A in various pathophysiological processes would help expand our current understanding of the biology of this extracellular lipolytic enzyme family. Lastly, in terms of human relevance, it is important to obtain more solid evidence that the PLA2G12A-driven lipid pathway is also operative in human Th17 cells with a view to future application of an anti-PLA2G12A antibody or a small-molecule PLA2G12A inhibitor to human diseases.

## MATERIALS AND METHODS

### Mice

*Pla2g12a*^-/-^ (TF0979; Taconic) and *Pla2g12a*^+/+^ mice on the C57BL/6 and DBA/1 backgrounds were obtained by intercrossing male and female *Pla2g12a*^+/–^ mice. *Lpar1*^-/-^ ^103^, *Lpar2*^-/-^ ^104^, and *Lpar6*^-/-^ ^105^ have been described previously. *Pla2g12a*^fl/fl^ mice ^29^ were crossed with *Cd4*^cre^ (B6-Tg(Cd4-cre)1Cwi/BfluJ; JAX:022071; The Jackson Laboratory) and *Cola2*^cre^ (C57BL/6-Tg(Col1a2-Cre)Kyo; nbio228; Laboratory Animal Resource Bank, National Institutes of Biomedical Innovation, Health and Nutrition) mice to obtain CD4^+^ T cell- and fibroblast-specific *Pla2g12a*-deficient mice, respectively. Mice were housed in climate-controlled (23 ± 1 ℃) and humid-controlled (50 ± 10%) specific pathogen-free facilities with a 12-h light-dark cycle, with free access to standard laboratory food (CE2 Laboratory Diet, CLEA Japan) and ultrafiltration water. Male 8- to 15-week-old mice were used for all animal experiments. All animal experiments were performed in accordance with protocols approved by the Institutional Animal Care and Use Committees of the University of Tokyo in accordance with the Japanese Guide for the Care and Use of Laboratory Animals (approval numbers; M-P17-032 and M-P20-102).

### Mouse genotyping

Mice were identified by ear punching. Tails were cut 3-5 mm, added with 30 μL of All^ele^– In–One–Mouse Tail Direct Lysis Buffer (Allele Biotech), and incubated overnight at 55°C with shaking. Mouse genotypes were determined by PCR of tail-snip DNA using GeneAmp^®^ Fast PCR Master Mix (Thermo Fischer Scientific-Applied Biosystems) and genotyping primers (Fasmac) on an Applied Biosystems 9800 Fast Thermal Cycler (Thermo Fischer Scientific-Applied Biosystems). The primer sequences used were as follows: wild-type (WT) forward 5’-GAAGCGGGAAGGACGAG-3’ and WT reverse 5’-CATGACCGCACGTGCTAAACTC-3’ for *Pla2g12a* WT allele (PCR product size, 328 base pairs); knockout (KO) forward 5’-GCAGCGCATCGCCTTCTATC-3’ and KO reverse 5’-TTCTCATGGGAACAACATAGCTC-3’ for the *Pla2g12a* mutant allele (PCR product size, 411 base pairs).

### Isolation of CD4^+^CD62L^+^ naïve T cells and ex vivo Th0/Th17 culture

*Ex vivo* Th0/Th17 differentiation culture was described previously ^29^. Mouse spleens were harvested and treated with 2 ml of ACK lysing buffer (Thermo Fischer Scientific-Gibco) at room temperature for 2 min, passed through 40-µm filters (Greiner Bio-One), centrifuged at 300 × *g* for 5 min, and resuspended in FACS buffer containing phosphate-buffered saline (PBS; Fujifilm Wako), 3% (*v*/*v*) fetal bovine serum (FBS, Biosera), and 2 mM EDTA (Dojindo). The cells were counted with a TC20 Automated Cell Counter (Bio-Rad). Mouse CD4^+^CD62L^+^ naïve T cells were isolated using a MojoSort Mouse CD4 T Cell Isolation Kit (BioLegend) on an autoMACS Pro Separator (Miltenyi Biotec). Naïve T cells (10^6^ cells/ml) were plated onto 24-, 48-, or 96-well plates precoated overnight at 4°C with 2 µg/ml anti-CD3ε antibody (145-2C11, BioLegend). For Th0 and Th17 differentiation, the cells were cultured with (for Th17) or without (for Th0) 10 ng/ml recombinant mouse IL-1β, 10 ng/ml mouse IL-6, 10 ng/ml mouse IL-23, and 2 ng/ml mouse TGF-β in the presence of 2 µg/ml anti-IFN-γ antibody (XMG1.2), 2 µg/ml anti-IL-4 antibody (11B11), and 2 µg/ml anti-CD28 antibody (37.51) (all from BioLegend) in RPMI1640 medium (Fujifilm Wako) containing 1 mM sodium pyruvate, 50 µM 2-mercaptoethanol, 25 mM HEPES, 1% (*v*/*v*) non-essential amino acids, and 10% (*v*/*v*) FBS for up to 3 days at 37°C under 5% CO_2_. As required for experiments, Th17-derived EVs (100 µg/ml protein equivalents; see below) pretreated for 4 h with or without 1 μg/ml PLA2G12A (provided by Dr. G. Lambeau), 0.1–1 µM lysophospholipids (LPC 16:0, LPE 18:0, LPE18:1, LPA16:0 or LPA18:0; all from Avanti Polar Lipids), 1 µM DBIBB (LPA_2_ agonist; Cayman Chemical) ^50^ or other LPA receptor agonists (T16, T17, T13, T7, T8, T10, and T19) ^49^, 1 µM PF-8380 (ATX inhibitor; Cayman Chemical) ^106^, 0.1–10 µM Ki16425 (LPA_1/3_ antagonist; Cayman Chemical) ^107^, 10 µM Y-27632 (ROCK1/2 inhibitor, Cayman Chemical),10 µM KD025 (ROCK2 inhibitor, Cayman Chemical), 10 µM YM254890 (G_q_ inhibitor, Fujifilm Wako), 10 µM NF-499 (G_s_ inhibitor, Cayman Chemical), or 10 µM pertussis toxin (PTX; G_i_ inhibitor; Vendors) were added to the Th17 culture.

### qPCR

Total RNA was isolated from cells or tissue using TRIzol reagent (Thermo Fisher Scientific-Invitrogen) and reverse transcription was carried out using the High-Capacity cDNA Reverse Transcription Kit (Thermo Fisher Scientific-Applied Biosystems). qPCR was performed using TaqMan Gene Expression Master Mix or PrimeTime Gene Expression Master Mix and TaqMan probe/primer sets (TaqMan Gene Expression Assay, Applied Biosystems; PrimeTime qPCR Probe Assays, and Integrated DNA Technologies) listed in Table S1 on the StepOnePlus Real-Time PCR System (Thermo Fisher Scientific-Applied Biosystems). The mRNA expression level was calculated as the ΔCt value (Ct value of target gene - Ct value of endogenous control gene), and the 2^-ΔCt^ was used as the relative gene expression level. The fold changes were also calculated using the ΔΔCt method.

### Microarray analysis

Agilent Technologies instruments and reagents were used for all analyses. Total RNA was extracted from Th17 and Th0 cells, and the RNA quality was assessed using the 2100 Bioanalyzer. The reverse transcription and *in vitro* transcription Cy3 labeling processes were carried out using the Low Input QuickAmp Labelling Kit. The prepared cRNA was hybridized overnight with an oligo DNA array (Whole Mouse or Human Genome DNA Microarray Kit, 4 x 44 K) carrying approximately 44,000 genes. After washing, the array was scanned using the SureScan Microarray Scanner, and the fluorescence intensity of each spot was converted into data using the Feature Extraction software. GeneSpring GX software was used to standardize raw data, compare gene expression between groups, and perform gene ontology analysis.

### Psoriasis model

The IMQ-induced psoriasis model was performed as described previously ^86,108,109^. IMQ (BESELNA cream 5%, Mochida Pharmaceutical Co., Ltd) was applied topically to mouse skin once a day for 6 days. The ear thickness was repeatedly measured with a micrometer (MDQ-30; Mitsutoyo). The ear skins, draining LNs, and spleens were then collected and subjected to flow cytometry and qPCR. For histological analysis, mouse ears were fixed in 10% (*v*/*v*) formalin neutral buffer solution (Fujifilm Wako) overnight at 4°C. The fixed tissue was placed in Unicaset (Sakura), dehydrated by immersion in 70% (*v*/*v*) ethanol, and then the samples were placed in a basket and replaced with paraffin using Tissue-Tek VIP-5-Jr. (Sakura). The samples were embedded in paraffin, heated to 60°C using a Tissue-Tek TEC (Sakura), and solidified at 4°C. The embedded samples were placed on a RETORATOME (REM-710; Yamato) and cut into 4-µm thick sections. The samples were immersed in a 45°C water bath, adhered to a glass slide, dried at 45°C overnight using a Slide Warmer (Sakura), and then placed in a staining basket for deparaffinization. After washing with running water, the sections were stained with hematoxylin and eosin (HE). After dehydration and clearing, the sections were mounted in Softmounts (Wako) and observed using an All-in-One microscope (BZ-X710; Keyence).

### Arthritis models

The CIA model was described previously ^38,110,111^. Briefly, the immunization grade chicken type II collagen (Chondrex) was gently mixed with an equal volume of complete Freund’s adjuvant (CFA containing *M. tuberculosis*; Chondrex) according to the manufacturer’s instructions. *Pla2g12a*^+/+^ and *Pla2g12a*^-/-^ mice on the DBA/1 background were anesthetized and 100 µl (100 µg collagen per mouse) of an emulsion of type II collagen and CFA containing *M. tuberculosis* at a final concentration of 2 mg/ml was injected intradermally at multiple sites at the base of the tail. 21 days after the primary immunization, the mice were received a booster injection of the same concentration of type II collagen in emulsion with incomplete Freund’s adjuvant (Chondrex). The mice was monitored by swelling of each paw and was scored for arthritis severity (0 = normal; 1 = slight swelling and/or erythema; 2 = pronounced swelling; 3 = ankyloses), as described previously ^111^. For µCT analysis, femurs preserved in 70% (*v*/*v*) ethanol were scanned using a microfocus X-ray CT system (inspXio SMX-90CT, Shimadzu). The µCT scanning conditions were as follows: tube voltage 90 kV; tube current 120 μA; and TRI/3D-BON (Ratoc) was used for imaging analysis. For histological analysis, ankle tissues were soaked in 4% (*v*/*v*) paraformaldehyde phosphate buffer solution (Fujifilm Wako) overnight and then replaced with OSTEOSOFT (Merck Millipore) every three days over 11 days for decalcification. The tissue sections were stained using the TRAP/Alkaline phosphatase Stain Kit (Fujifilm Wako) according to the manufacturer’s instructions.

For the K/BxN serum-induced arthritis, the K/BxN mouse serum (provided by Dr. E. Boilard) was intraperitoneally injected into *Pla2g12*^+/+^ and *Pla2g12a*^-/-^ mice on the C57BL/6 background on days 0 and 2, as described previously ^112,113^. The ankle thickness was measured using a micrometer at various times before and after onset of the disease. The ankle tissues on day 4, a peak time of tissue swelling, were subjected to qPCR analysis.

### BM transplantation

*Pla2g12a*^-/-^, *Lpar1*^-/-^, and their control WT mice were used as donors or recipients. The recipients were irradiated with a lethal dose (10.4 Gy) using an X-irradiation system (M-150WE, Softex) equipped with a RAMTEC Solo Dosimeter (Ramtec), and then 10^7^ BM cells from the donors were injected intravenously into the recipients. Eight weeks later, IMQ-induced psoriasis was applied to the recipients.

### Dispersion of skin tissue

The ear skin was minced finely with scissors and digested in RPMI1640 medium containing 333-666 µg/ml Liberase TL (1 vial with 5–10 mg pack size; Roche), 2 kU/ml DNase I (Merk Sigma-Aldrich), and 10% (*v*/*v*) FBS at 37°C for 1 h, and then treated with 10 mM EDTA at 37°C for 5 min. After washing, the resultant cells were isolated by 25% (*v*/*v*) Percoll (Cytiva) density gradient, and then filtered through a 70-µm cell strainer. The cells were resuspended in FACS buffer and quantified by flow cytometry, as described below.

### Flow cytometry (FACS)

For intracellular cytokine staining, the cells were stimulated with 500 nM ionomycin and 10 ng/ml phorbol-12-myristate-13-acetate (PMA) (both from Merck Sigma-Aldrich), and BD GolgiPlug Protein Transport Inhibitor (containing Brefeldin A) (Fisher Scientific). The cells were incubated with TruStain FcX Fc Receptor Blocking Solution (anti-mouse CD16/32 antibody; BioLegend) to block nonspecific binding to Fc receptors for 10 min on ice, and then centrifuged at 500 × *g* for 5 min at 4°C. The cells were resuspended in FACS buffer and incubated with fluorescence-labeled antibodies against cell surface markers (listed in Table S2) for 30 min in the dark on ice. After washing, the cells were incubated in 100 μl of Cytofix/Cytoperm (BD Biosciences) for 20 min in the dark on ice, washed with 1x Perm/Wash Buffer (BD Biosciences), and then incubated with PE-labelled anti-mouse IL-17A antibody (eBiosciences) at 1:100 dilution for 30 min in the dark on ice. After washing, the cells were resuspended in 500 μl of FACS buffer, passed through 40-μm cell strainers, and analyzed using a BD FACSMelody Cell Sorter (BD Biosciences) and Flowjo 10.9.0 software.

### LC-ESI-MS/MS

Lipidomics analysis was performed by LC-ESI-MS/MS system according to our previous protocols ^41,42^. In brief, for analysis of phospholipids and lysophospholipids, lipids were extracted using the Bligh and Dyer method ^114^. MS analysis was performed on a triple quadrupole linear ion trap mass spectrometer QTRAP4000 or QTRAP4500 (ABSciex) with Nexera UPLC system (Shimadzu).

Quantification of phospholipids and lysophospholipids were performed on the negative ion mode with multiple reaction monitoring (MRM) transition using QTRAP4000. Alternatively, neutral and acidic phospholipids or lysophospholipids were quantified on positive and negative ion modes, respectively, with MRM transitions using QTRAP4500. When using a reversed-phase column, the samples were separated with a Kinetex C18 column (1.7 µm, 150 × 2.1 mm, Phenomenex) maintained at 50°C using mobile phase A [acetonitrile/methanol/water = 1:1:1 (*v*/*v*/*v*) containing 5 µM phosphoric acid (Fujifilm Wako) and 1 mM ammonium formate] and mobilephase B [2-propanol containing 5 µM phosphoric acid and 1 mM ammonium formate] with a flow rate of 0.2 mL/min. When using the hydrophilic column, the samples were separated with a SeQuant ZIC-HILIC column (2.1 x 250 mm; 3.5 µm), eluted by a mixture of mobilephase A (acetonitrile/water = 95/5 (*v*/*v*), 10 mM ammonium acetate) and mobile phase B (acetonitrile/water = 50/50, 20 mM ammonium acetate) at 40°C with flow rate of 0.3 ml/min.

For detection of fatty acid metabolites, tissues were soaked in methanol and homogenized in a bead homogenizer. After incubation at -20°C overnight, water was added to bring the final methanol concentration to 10% (*v*/*v*). samples in 10% methanol were applied to an Oasis HLB cartridge (Waters), washed with 10 mL hexane, eluted with 3 mL methyl formate, dried under N_2_ gas, and dissolved in 60% (*v*/*v*) methanol. Quantification of fatty acid metabolites were performed on the negative ion mode with MRM transition using QTRAP4000. The samples applied to Kinetex C18 column (1.7 µm, 150 × 2.1 mm, Phenomenex) was separated using a linear gradient with mobile phase C [water containing 10 mM ammonium acetate (Fujifilm Wako)] and mobile phase D [acetonitrile/methanol = 4:1 (*v*/*v*)] at a flow rate of 0.2 ml/min at 40°C. As internal standards, a cocktail consisting of phospholipid [PC, PE, PGl, PI and PS with 12:0 and 13:0 acyl chains at the *sn*-1 and *sn*-2 positions, respectively, PA with 17:0–17:0, plasmalogen-type PE with 18:1–18:1 (*d*9); Avanti Polar Lipids], and lysophospholipids [LPC with 16:0 (*d*49), LPE with 18:1 (*d*7), LPA with 17:0, LPG, LPI, and LPS with 17:1; all from Avanti Polar Lipids] was added to each sample. For data analysis, peak area values for each molecule were obtained using MultiQuant software (ABSciex), and quantification was performed based on the peak area of the MRM transition and the calibration curve obtained with an authentic standard for each compound [LPC, LPE, P-LPE (lysoplasmalogen), and LPS with 18:1, LPA and LPG with 16:0, and LPI with 18:0 (all from Avanti Polar Lipids)].

### Isolation of EVs

Th17 cells (10^7^ cells/ml) were cultured for 2 days in the Th17 differentiation medium containing 10% exosome-depleted FBS (Biosera). The culture supernatant was centrifuged at 500 x *g* for 10 min to remove the cell fraction, and then microfiltered using a 0.22-μm syringe filter (Merck Millipore). EVs in the culture supernatant were concentrated in an ultrafiltration tube (Vivaspin 20, Sartorus) at 100,000 x *g* for 70 min at 4°C using the ultracentrifuge Optima MAX-TL (Beckman Coulter). The pellets were suspended in PBS(–) and ultracentrifuged again. The isolated EVs were resuspended in 50 μl of PBS(–), quantified their protein contents using the Pierce BCA Protein Assay Kit (Thermo Fisher Scientific), and used for experiments. To evaluate CD9^+^CD69^+^ EVs, they were captured with a combination of anti-CD9-labeled discs and anti-CD63-labeled nanobeads and counted with an EXOCOUNTER (JVCKENWOOD).

### TEM

Carbon film grids (ELS-C10; STEM) were hydrophilized using PIB-10 (Shinku Device) under soft conditions for 1.5 min, and then 5 mg/ml EVs were applied and excess solution was removed with filter papers. Grids were stained with 1% (w/v) uranyl acetate solution on the JEM-1400Flash electron microscope (JEOL) at 100 kV.

### Cryo-EM

QUANTIFOIL (QUANTIFOIL 1.2/1.3, 300 mesh, Au; Quantifoil) was hydrophilized by PIB-10 under hard conditions for 2 min and then coated with 5 mg/ml EVs. Grids were blotted using a Vitrobot Mark IV system (FEI; ThermoFisher Scientific) at 6°C under 100% humidity, with blotting time of 4 seconds, and immersed in liquid ethane. The prepared grids were imaged using a Talos Arctica TEM (ThermoFisher Scientific) operated at 200 kV and a Falcon 3. Data were collected using an EPU.

### Immunofluorescence microscopy of EV uptake

EVs were stained with ExoSparkler Exosome Membrane Labeling Kit Red (Dojindo). T cells (5 x 10^4^ cells) were seeded in 96-well plates and cultured at 37°C for 2 days. After washing with PBS, 10 μg of labeled EVs were added and cultured at 37°C for 4 h. Cells were sealed with DAPI and observed with an All-in-One microscope.

### Small RNA sequencing

RNA was extracted from EVs purified using microRNA ExtractorR Kit for Purified EV (Fujifilm Wako) and adjusted at 15 ng/µl in RNase-free water according to the manufacturer’s instructions. The Small RNA sequencing library was independently prepared by RIKEN IMS (Japan) using the NEBNext Multiplex Small RNA Library Prep Set for Illumina (New England Biolabs) with 200 ng of total RNA for each sample. The libraries were submitted to Illumina NextSeq 2000 (Illumina), and paired-end sequencing (2 x 50 bp) was performed at RIKEN IMS.

Fastq files were generated using bcl2fastq software (version 2.20.0.422) with base mask option Y51,I6,Y51 (51bp paired-end with 6bp barcode). QC and trimming were performed using by trim galore (version 0.6.7). Mapping with STAR (version 2.5.4) and count estimation with RSEM (version v1.3.0) were performed. Annotation is provided by Gencode (gencode.vM25.annotation.gtf). Within each sample, expression levels for mature microRNAs were normalized to reads per million mapped reads (RPM).

### Proteomics analysis

Shotgun proteomic analysis was performed as described previously, ^115^ with modifications. The EV samples (5 µg protein equivalents) dried in PCR tubes were added to 25 µl of methanol. The samples were sonicated for 3 min using a sonicator (UIP400MTP, Hielscher) and then centrifuged at 4°C for 20 min at 19,000 *g*. After removal of the supernatant (20 µl), the samples were centrifuged to dryness. The proteins were dissolved by sonication in 4 µl of trypsin/Lys-C solution (100 ng/µl) in 50 mM ammonium bicarbonate. The mixture was incubated at 37°C for 2 h while shaken with a Thermomixer comfort (Eppendorf) at a setting of 300 rpm, diluted with 1 µl of 1.25% (*v*/*v*) trifluoroacetic acid (TFA), and subjected to nano-flow liquid chromatography high-resolution tandem mass spectrometry (nano-LC/MS/MS) analysis.

The nano-LC/MS/MS system was composed of a Dionex Ultimate 3000 nano-RSLC system and a Q-Exactive HF, a high-performance benchtop quadrupole Orbitrap mass spectrometer (Thermo Fisher Scientific) equipped with a Dream spray electrospray ionization source (AMR Inc). An Acclaim™ PepMap™ C18 column (Thermo Fisher Scientific) with dimensions of 300 µm inner diameter (i.d.) × 5 mm and a particle size of 5 µm was used as the pre-column for sample trapping. The loading pump was run at 1 µl/min with water/acetonitrile/TFA (98/2/0.1, *v*/*v*/*v*) and 1 µl was injected per sample. After loading, the sample was switched online to the packed nano-LC column of 2 µm particle-size L-column2 ODS (CERI, Saitama, Japan) with dimensions of 50 µm i.d. × 200 mm. The nano-LC conditions were as follows: mobile phase, water/formic acid (100/0.1, v/v) (solvent A) and water/acetonitrile/formic acid (80/20/0.1, v/v/v) (solvent B); flow rate, 200 nl/min; and column temperature, 40°C. The gradient conditions were as follows: 5% B, 0−3 min; 5%−35 B, 3−63 min; 35%−80% B, 63−64 min; 80% B, 64−69 min; 80%−5% B, 69−70 min; and 5% B, 70−90 min.

The MS analysis conditions were as follows: polarity, positive ionization; spray voltage, 1.8 kV; capillary temperature, 275°C; S-lens level, 50; probe heater temperature, 350°C; mass resolution, 35,000; automatic gain control (AGC) target (the number of ions used to fill the C-trap), 1,000,000; maximum injection time (IT), 50 ms; and MS scan range, 430–860 (*m/z*). The MS/MS spectra were acquired using data-independent acquisition (DIA) mode with higher-energy collision dissociation. The DIA conditions were as follows: mass resolution, 35,000; AGC target, 200,000; maximum IT, auto; loop count, 21; isolation window, 20 Da; fixed first mass, 200 (*m/z*); and normalized collision energy (NCE), 25 eV.

Raw files from DIA were analyzed in DIA-NN 1.8 ^116^ using an *in silico* DIA-NN predicted spectral library. The parameters of the mouse reference spectral library generated from the SwissProt database (2023.03.18-21.42.04.29) are as follows: enzyme used, trypsin; peptide length range, 7−50; allowed number of maximum missed cleavages, 1; modifications, methionine oxidation, and *N*-terminal acetylation; and false discovery rate (FDR), less than 1%.

### Generation of anti-PLA2G12A monoclonal antibodies

The gene encoding human PLA2G12A (residues 23 to 189) was cloned from full-length human *PLA2G12A* cDNA (provided by Dr. G. Lambeau) in the modified pcDNA3.4 vector (Thermo Fisher Scientific) harboring an *N*-terminal human IL-2 signal sequence and *C*-terminal mIgG2aFc (Fc of mouse IgG2a), tobacco etch virus (TEV) protease cleavage site, and hexa histidine (His_6_)-tag. The recombinant PLA2G12-Fc fusion protein was transiently expressed using the Expi293 Expression System (Thermo Fisher Scientific) according to the manufacturer’s instructions. Culture media were harvested 7 days after transfection and the secreted PLA2G12A protein was purified from the culture supernatants by affinity chromatography with a HisTrap HP column (Cytiva). Then the His_6_-tag was cleaved by TEV protease at 4°C overnight. The protein was reloaded on a HisTrap HP column to remove the TEV protease and uncleaved protein. The flow-through fractions were further purified by size-exclusion chromatography on a HiLoad 16/60 Superdex 200 column (Cytiva) equilibrated in PBS. Peak fractions were concentrated using Amicon Ultra spin concentrator (Merck). The purified PLA2G12A protein was immunized into *Pla2g12a*^-/-^ mice by an iliac LN method ^117,118^ (ITM Inc). Hybridoma clones producing IgG_1_ antibodies that showed high reactivity to both human and mouse PLA2G12A proteins were screened by ELISA using their culture supernatants.

### PLA_2_ enzyme assay

PLA_2_ enzymatic activity was measured using a PLA_2_ Enzyme Assay Kit (Cayman) according to the manufacture’s instruction. Human or mouse recombinant PLA2G12A proteins (50 ng) was incubated for 4 h at room temperature with substrate phospholipids (diheptanoyl-thio-PC) in the presence or absence of various concentrations of anti-PLA2G12A antibodies. Alternatively, phospholipids extracted from mouse skin were incubated with various concentrations of human PLA2G12A for 4 h at 37°C, and the amounts of free fatty acids and lysophospholipids were quantified by LC-ESI-MS/MS, as described previously ^41,42^.

### Immunohistochemistry

Paraffin-embedded sections were deparaffinized and antigen was activated with 20 μg/ml proteinase K (Merk Sigma-Aldrich). Endogenous peroxidase was inactivated by treatment with 3% (*w*/*v*) hydrogen peroxide solution (Fujifilm-Wako) for 5 min, and then blocked with Block Ace (DS Pharma Biomedical) for 1 h. The sections were incubated with anti-PLA2G12A antibody (clone #44) biotinylated with a Biotin-Labeling Kit (Dojindo) and rabbit anti-CD4 monoclonal antibody (D7D2Z; Cell Signaling) diluted at 1:1000 at room temperature for 1 h. After washing, the sections were incubated with APC streptavidin (BioLegend) and Alexa Flour 488-conjugated F(ab’)_2_-goat anti-rabbit IgG (H+L) cross-adsorbed secondary antibody (A-11070, Thermo Fisher Scientific) diluted at 1:1000 at room temperature for 1 h. After washing, the sections were sealed with VECTASHIELD mounting medium with 4’,6-diamidino-2-phenylindole (DAPI) (Vector Laboratories).

### Western blotting

RIPA Buffer (Nacalai Tesque) mixed with protease inhibitors (cOmplete Protease Inhibitor Cocktail tablets) and phosphatase inhibitors (PhosSTOP Phosphatase Inhibitor Cocktail tablets) (both from Roche) was added to cell lysates. The protein concentrations were determined using the BCA assay, and cell lysates (10 µg protein equivalents) or recombinant human and mouse PLA2G12A proteins and human PLA2G3 protein (sPLA_2_ domain-only form)^119^ (50 ng) were subjected to sodium dodecyl sulfate polyacrylamide gel electrophoresis (SDS-PAGE) using 10 or 15% gels under reducing conditions, and transferred to PVDF membranes (Immobilon-P, Merck Millipore) using a Semi-Dry and Rapid Blotting Systems (Bio-Rad). The membranes were blocked with Block Ace and then probed with biotinylated anti-human PLA2G12A monoclonal antibody (clone #44) or mouse monoclonal antibodies for STAT3 (124H6), phospho-STAT3 (Tyr705) (M9C6), and β-actin (8H10D10) (all from Cell Signaling) at 1:1000 dilution in Can Get Signal Solution 1 (Toyobo) overnight at 4°C. After washing with 1x TBST [50 mM Tris, 140 mM NaCl, 2.5 mM KCl, and 0.005% (*v/v*) Tween-20 (Fujifilm Wako)], the membranes were incubated with horseradish peroxidase (HRP)-conjugated streptavidin (BioLegend) for PLA2G12A or HRP-conjugated sheep anti-mouse IgG F(ab’)_2_ fragment (Amersham) for STAT3, pSTAT3, and β-actin in Can Get Signal Solution 2 (Toyobo) at 1:1000 dilution at room temperature for 1 h. The signal was visualized with the ECL (SuperSignal West Pico PLUS Chemiluminescent Substrate, Thermo Fisher Scientific) on a Chemiluminescence Imaging System (FUSION Solo S, Vilber-Smart Imaging).

### ELISA

The level of IL-17A in mouse serum was determined using a Mouse IL-17A ELISA Kit (Proteintech). For evaluation of the serum titer of anti-type II collagen antibody, ELISA plates (Corning) were coated overnight with ELISA Grade bovine type II collagen (Chondrex) diluted to 0.5 mg/ml with 10x Collagen Dilution Buffer (Chondrex), blocked, and then incubated with serum samples at 1:1000 dilution for 2 h and then with HRP-conjugated sheep anti-mouse IgG F(ab’)_2_ fragment at 1:5000 dilution for 30 min at room temperature. After washing, the samples were subjected to a color reaction using 3,3’,5,5’-tetramethylbenzidine (TMB; ATTO) in the dark for 30 min. Then, stop solution (ATTO) was added and absorbance was measured at 450 nm. For evaluation of the specificity of anti-PLA2G12A antibody, ELISA plates were coated with various sPLA_2_s (provided by Dr. G. Lambeau) at 0.5 μg/ml for 1 h, blocked with Block Ace for 2 h, incubated with anti-PLA2G12A antibody (clone #44) for 1 h and then with HRP-linked rat anti-mouse IgG_1_ secondary antibody (eBioscience) at 1:5000 dilution for 30 min, and colored with TMB.

### Evaluation of the therapeutic efficacy of antibodies

In the IMQ-induced psoriasis model, mouse anti-PLA2G12A monoclonal antibody (clone #44), InVivoMAb anti-mouse IL-17A antibody (clone 17F3; Bio X Cell), or GoInVivo purified mouse IgG_1_, κ isotype control antibody (BioLegend) was administrated intraperitoneally at 1 mg/kg every day from day 0 to day 5 of IMQ application (Figure 8A). In the CIA model, the above antibodies were each administrated intraperitoneally at 1 mg/kg two days before the primary immunization, and then every seven days from day 0 to day 42 (Figure 8H). In the *ex vivo* Th17 differentiation, splenic naïve CD4^+^ T cells were cultured for 3 days with or without several anti-PLA2G12A monoclonal antibody clones at 0.1 μg/ml to assess IL-17A expression by FACS.

### Statistical analysis

Sample sizes were chosen based on previous experience in our laboratory. The experiments were performed and analyzed in a non-randomized and non-blinded manner. Data are expressed as Mean ± s.e.m.. GraphPad Prism version 9 (GraphPad) was used for statistical analysis. Differences in the statistical analysis between the two groups were determined by an unpaired t-test or Mann-Whitney U test, depending on the variance. For statistical analysis of three or more groups, one-way ANOVA with a post hoc Tukey’s, Šidák, and Dunnet multiple comparisons test, Brown-Forsythe and Welch ANOVA with Dunnett’s T3 multiple comparisons test, Mixed-effects model with Sidak’s multiple comparisons test, two-way repeated measures ANOVA with a post hoc Šidák multiple comparisons test, and Kruskal-Wallis with a post hoc Dunn’s multiple comparisons test were performed as noted in the respective figure legends. *P* < 0.05 (*), *P* < 0.01 (**) were considered significant.

## ACKNOWLEDGMENTS

We thank Dr. Eric Boilard (Centre de Recherche du Centre Hospitalier Universitaire de Québec -Université Laval, Canada) for providing the serum of K/BxN mice, Dr. Gerard Lambeau (Centre National de la Recherche Scientifique, France) for providing recombinant human and mouse sPLA_2_ proteins and cDNAs, Drs. Yoichi Sakamaki, Toshie Furuya, and Masahide Kikkawa (The University of Tokyo, Japan) for assisting TEM and cryo-EM analyses, Dr. Yoshiaki Kitaura (The University of Tokyo, Japan) for assisting μCT analysis, and Dr. Takehisa Matsumoto and Mayumi Yonemochi (RIKEN, Japan) for assisting protein preparation. This work was supported by JSPS KAKENHI Grants-in-Aid for Scientific Research (S) JP20H05691 (to M.M.) and (B) JP23H02940 (to A.K.) from the Japan Society for the Promotion of Science (JSPS), AMED-CREST JP24gm1210013 (to M.M.) and P-PROMOTE JP24ama221216 (to M.M.) from the Japan Agency for Medical Research and Development (AMED), JST-CREST JPMJCR19H5 (to A.K.) from the Japan Science and Technology Agency (JST), JST SPRING JPMJSP2108, Ono Medical Research Foundation (to Y.T.), and Cayman Biomedical Research Institute (CABRI) (to M.M.). This work was also in part supported by P-PROMOTE Advanced Technical Support and Efficient Promotion Management JP22ama221001, AMED-BINDS (Basis for Supporting Innovative Drug Discovery and Life Science Research) JP23ama121002, JP23ama121018, and JP23ama121055, and the MEXT Cooperative Research Project Program, Medical Research Center Initiative for High Depth Omics, and Coalition of Universities for Research Excellence Program (CURE) JPMXP1323015486 for Medical Institute of Bioregulation, Kyushu University.

## AUTHOR CONTRIBUTIONS

C. Mochizuki, Y. Taketomi, and M. Murakami designed the study and wrote the manuscript. Y. Miki, Y. Nagasaki, and T. Ono assisted lipidomics, qPCR, and immunohistochemistry. A. Irie performed the analysis using K/BxN mice. Y. Nishito and Y. Taketomi performed microarray analysis. Y. Endo and T. Nakajima assisted the analysis of Th17 cell differentiation and *Pla2g12a*^fl/fl^*Cd4*^cre^ mice. Y. Tomabechi, K. Hanada and M. Shirouzu supported protein purification. T. Watanabe performed small RNA-seq. K. Hata, Y. Izumi, and T. Bamba performed proteomics analysis. K. Kudo and A. Kotani assisted EV analysis. J. Chun generated *Lpar1*^-/-^ mice. K. Kano and J. Aoki assisted the studies using *Lpar1*^-/-^, *Lpar2*^-/-^ and *Lpar6*^-/-^ mice and LPA agonists or antagonists. All authors have read and agreed to the published version of the manuscript.

## DECLARATION OF INTERESTS

J. Chun has an employment relationship with Neurocrine Biosciences, Inc. as a Distinguished Scholar, unrelated to the current manuscript. The remaining authors have no conflicting financial interests.

---

**Figure S1.**
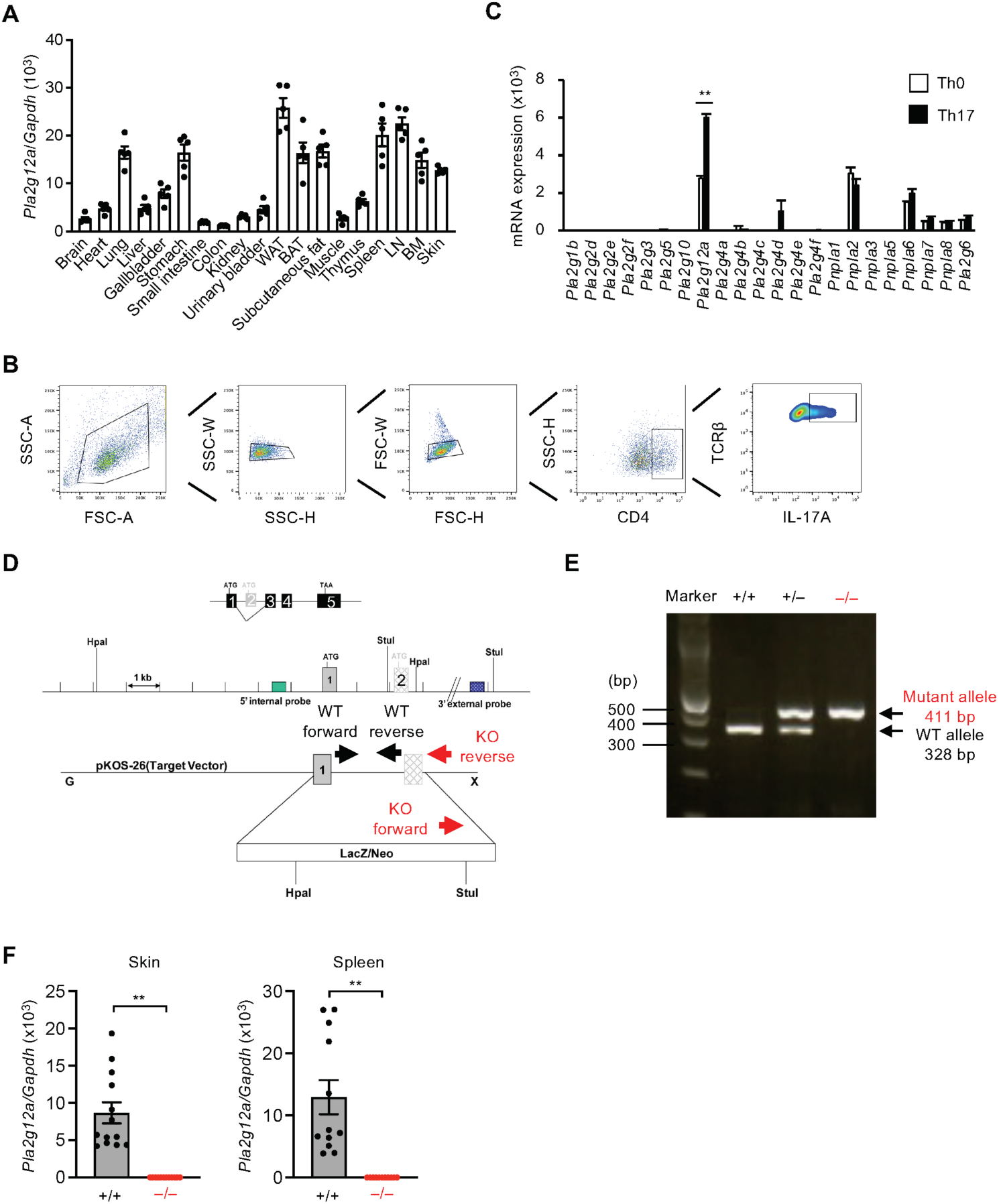
Generation of PLA2G12A-deficient mice, related to Figure 1. (A) qPCR of *Pla2g12a* in various tissues of 10-week-old C57BL/6 mice. (B) Procedure used for flow cytometry of CD4^+^-gated TCRβ^+^IL-17A^+^ Th17 cells from *ex vivo* Th17 differentiation culture. (C) qPCR of various PLA_2_ enzymes in splenic CD4^+^ T cells cultured for 3 days under Th0 and Th17 differentiation conditions (*n* = 3). (D) Strategy for gene targeting of *Pla2g12a*. Positions of PCR primers for genotyping are indicated by arrows. (E) PCR genotyping of *Pla2g12a*^+/+^ (+/+), *Pla2g12a*^+/–^ (+/–), and *Pla2g12a*^-/-^ (–/–) mice. (F) qPCR of *Pla2g12a* in the skin and spleen of *Pla2g12a*^+/+^ and *Pla2g12a*^-/-^ mice. Values are mean ± s.e.m.. Representative data of two experiments (C) and combined results of two experiments (A, F) are shown. Statistical analysis was performed by Mann-Whitney U test (B, F, G). **, *P* < 0.01.

**Figure S2.**
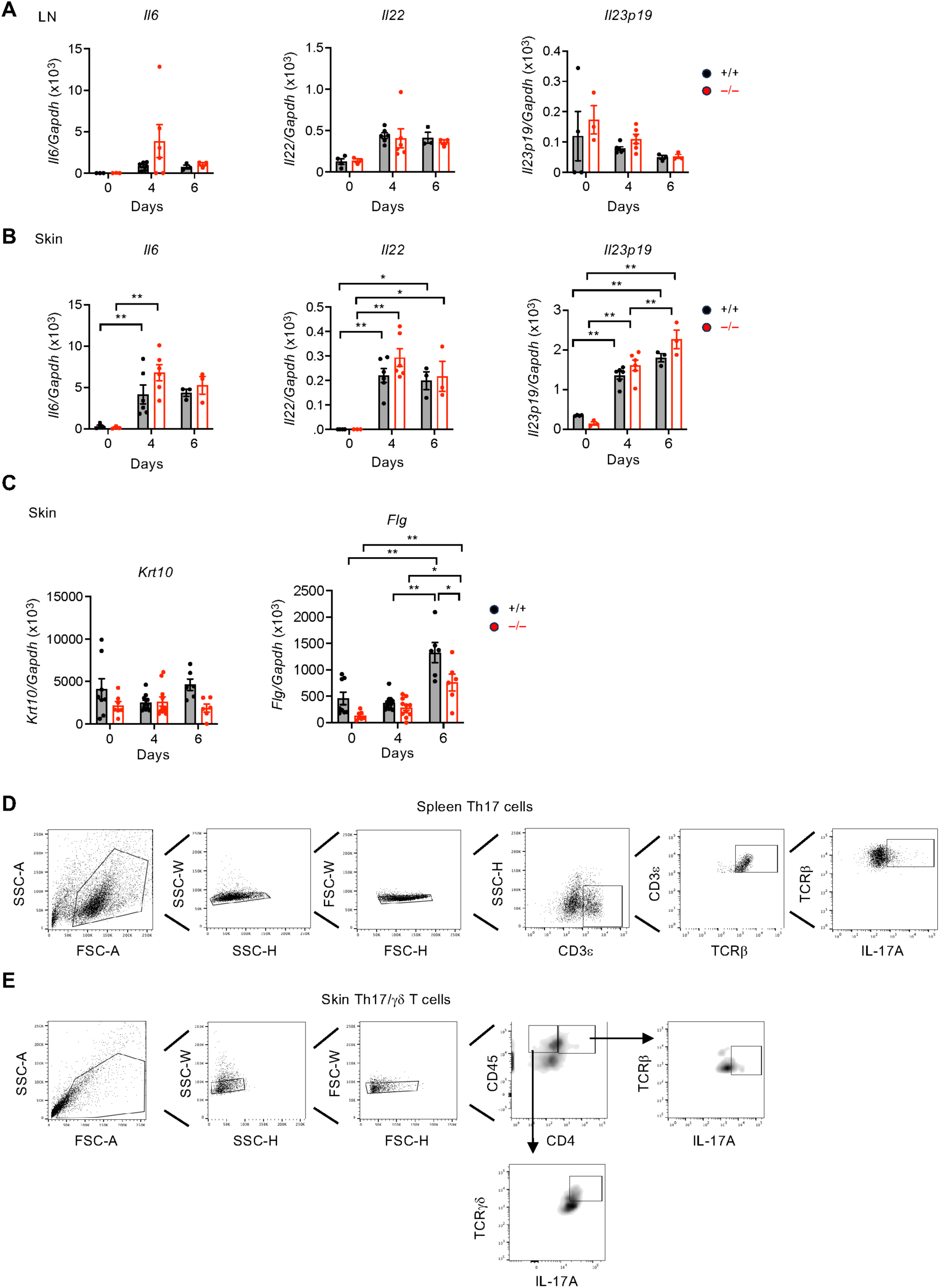
IMQ-induced psoriasis in *Pla2g12a*^+/+^ and *Pla2g12a*^-/-^ mice, related to Figure 2. (A-C) qPCR of cytokines (A, B) and keratinocyte markers (C) in the LNs (A) and skin (B, C) of *Pla2g12a*^+/+^ and *Pla2g12a*^-/-^ mice. (D, E) Procedures used for flow cytometry of CD3ε^+^-gated TCRβ^+^IL-17A^+^ Th17 cells from the spleen (D) and CD45^+^-gated TCRβ^+^IL-17A^+^ Th17 cells and TCRγδ^+^IL-17A^+^ γδ T cells from the skin (E). Values are mean ± s.e.m.. Results of one experiment (A, B) and combined results of two experiments (C) are shown. Statistical analysis was performed using two-way ANOVA with Sidak’s multiple comparisons test (A–C). *, *P* < 0.05; **, *P* < 0.01.

**Figure S3.**
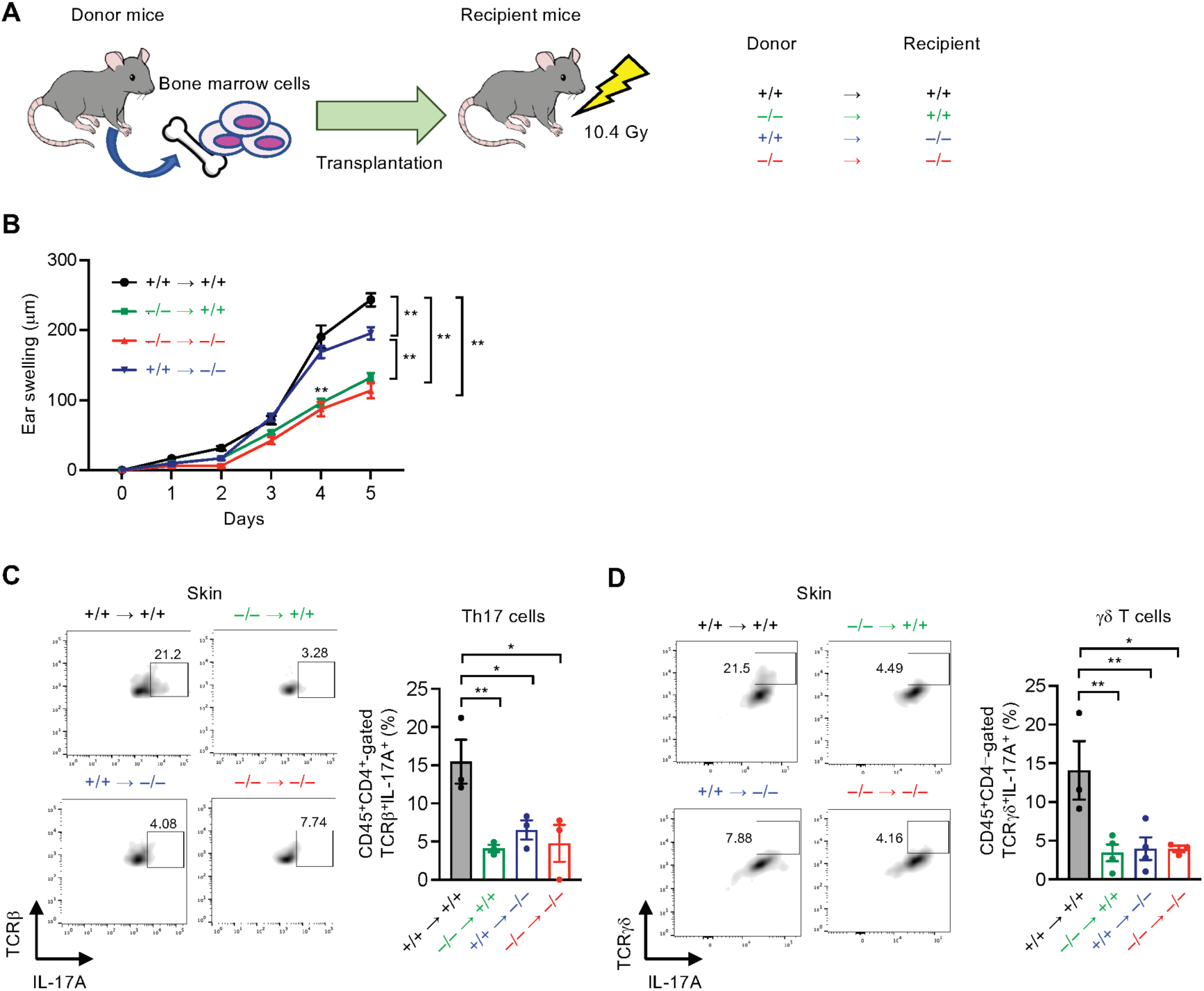
Adoptive transfer of *Pla2g12a*^+/+^ and *Pla2g12a*^-/-^ BM cells, related to Figure 3. (A) Schematic representation of the procedure used for BM transfer. (B) IMQ-induced psoriasis in *Pla2g12a*^+/+^ (+/+) and *Pla2g12a*^-/-^ (–/–) mice adoptively transferred with BM cells from *Pla2g12a*^+/+^ or *Pla2g12a*^-/-^ mice (*n* = 8-18). (C, D) FACS of cutaneous Th17 cells (C) and γδ T cells (D) on day 5 in (B). Representative FACS profiles (*left*) and the proportion and number of Th17 (C) or γδ T (D) cells (*right*) are shown. Values are mean ± s.e.m.. Representative data of two experiments (B) and results of one experiment (C, D) are shown. Statistical analysis was performed using one-way ANOVA Dunnett’s multiple test (C, D) and two-way repeated measures ANOVA with Sidak’s multiple comparisons test (B). *, *P* < 0.05; **, *P* < 0.01.

**Figure S4.**
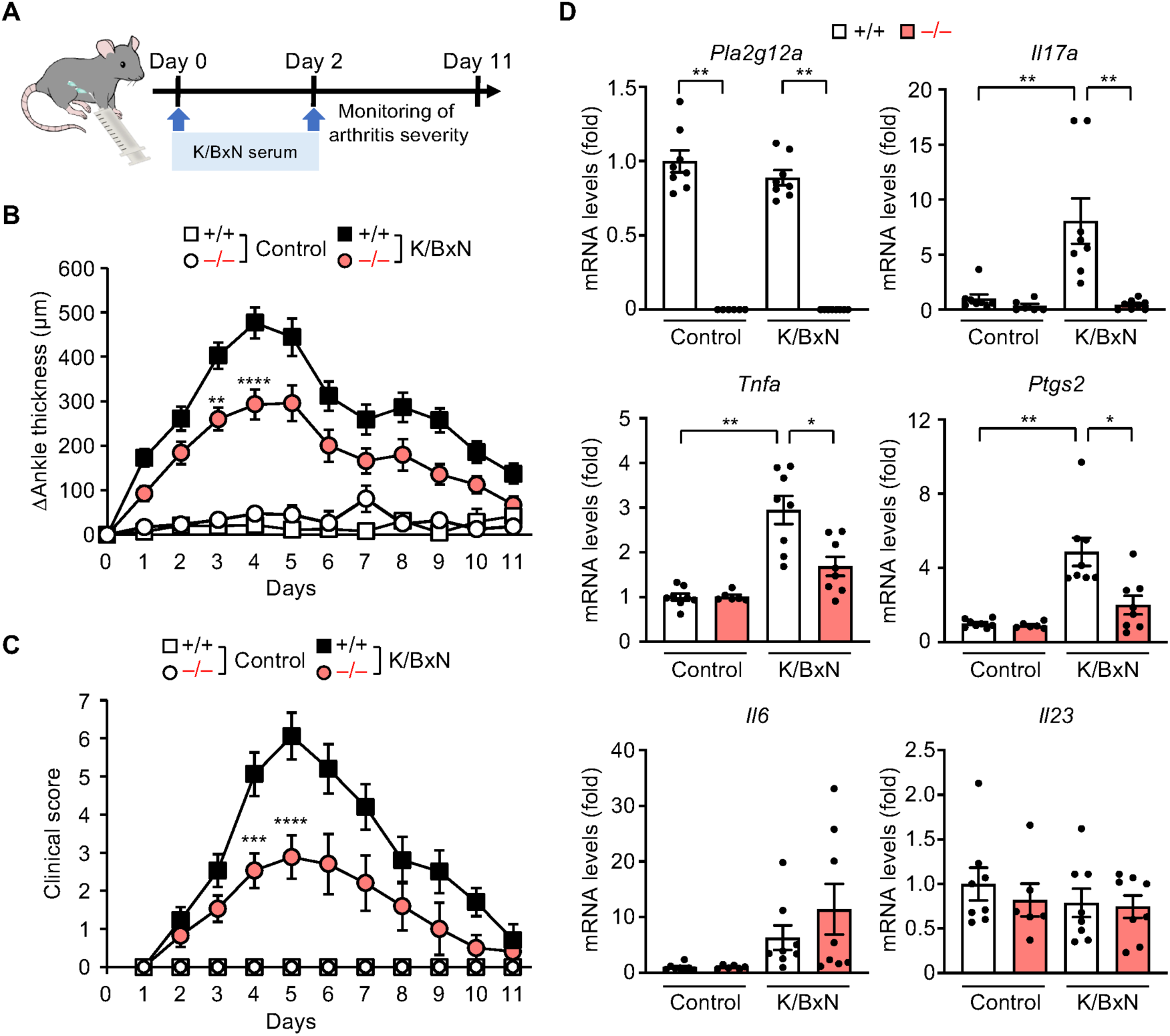
K/BxN serum transfer arthritis in *Pla2g12a*^+/+^ and *Pla2g12a*^-/-^ DBA/1 mice, related to Figure 4. (A) Schematic representation of the procedure used for K/BxN serum transfer arthritis. (B, C) Ankle swelling (B) and arthritis score (C) in *Pla2g12a*^+/+^ (+/+) and *Pla2g12a*^-/-^ (–/–) mice over 11 days. (D) qPCR of *Pla2g12a* and arthritis-related genes in the joints of *Pla2g12a*^+/+^ and *Pla2g12a*^-/-^ mice on day 4 in (B, C). Values are mean ± s.e.m. Combined results of two experiments are shown (B–D). Statistical analysis was performed using Brown-Forsythe and Welch ANOVA with Dunnett’s T3 multiple comparisons test (D) and two-way ANOVA with Tukey’s multiple comparisons test (B, C). *, *P* < 0.05; **, *P* < 0.01.

**Figure S5.**
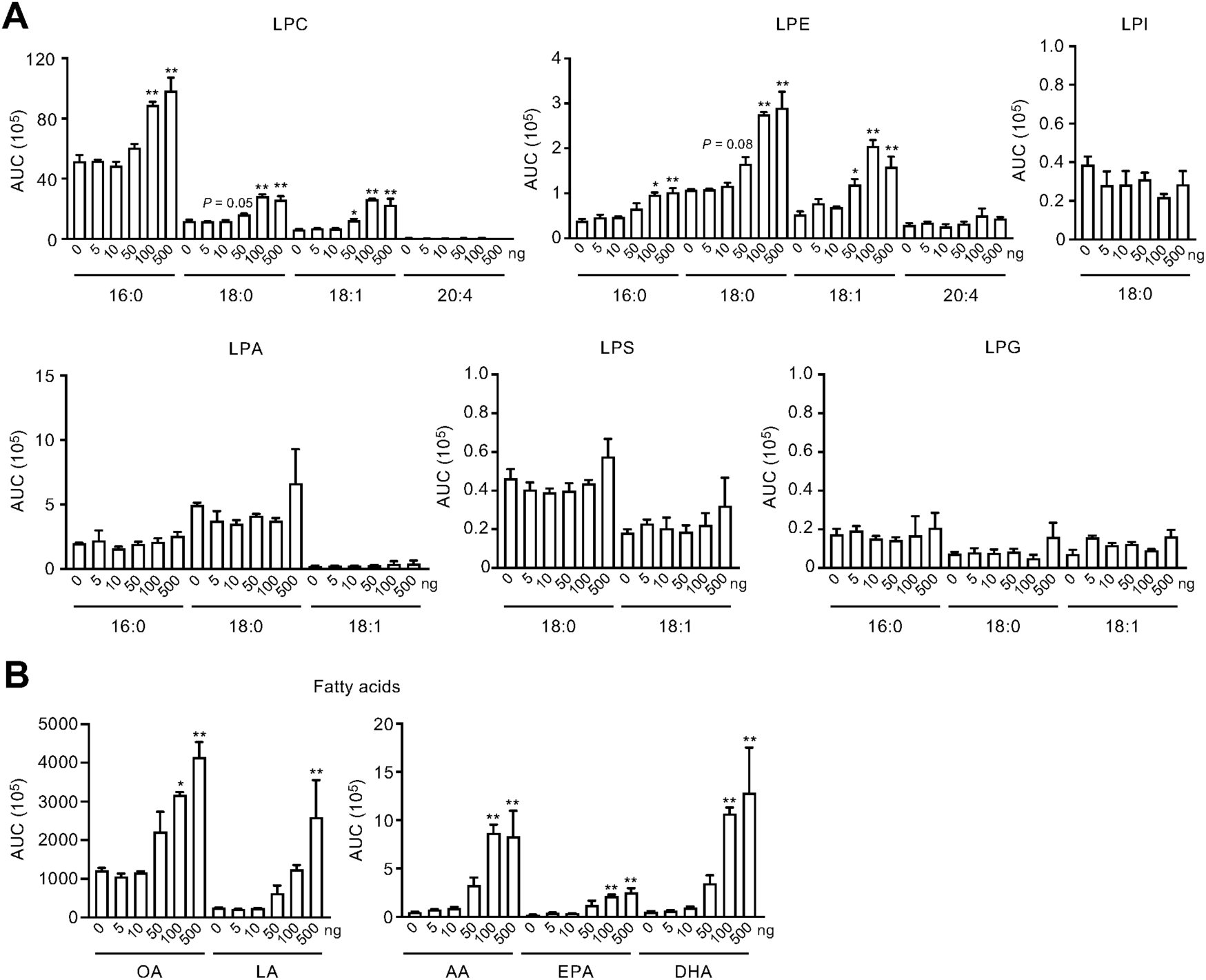
*In vitro* enzymatic activity of PLA2G12A, related to Figure 5. Skin-extracted phospholipids were incubated with various concentrations of recombinant human PLA2G12A for 30 min and the generation of lysophospholipids (A) and free fatty acids (B) was analyzed by LC-ESI-MS/MS (*n* = 6). Values are mean ± s.e.m.. Combined results of two experiments are shown. Statistical analysis was performed using ordinary one-way ANOVA with Dunnett’s multiple comparisons test. *, *P* < 0.05; **, *P* < 0.01 *versus* without PLA2G12A.

**Figure S6.**
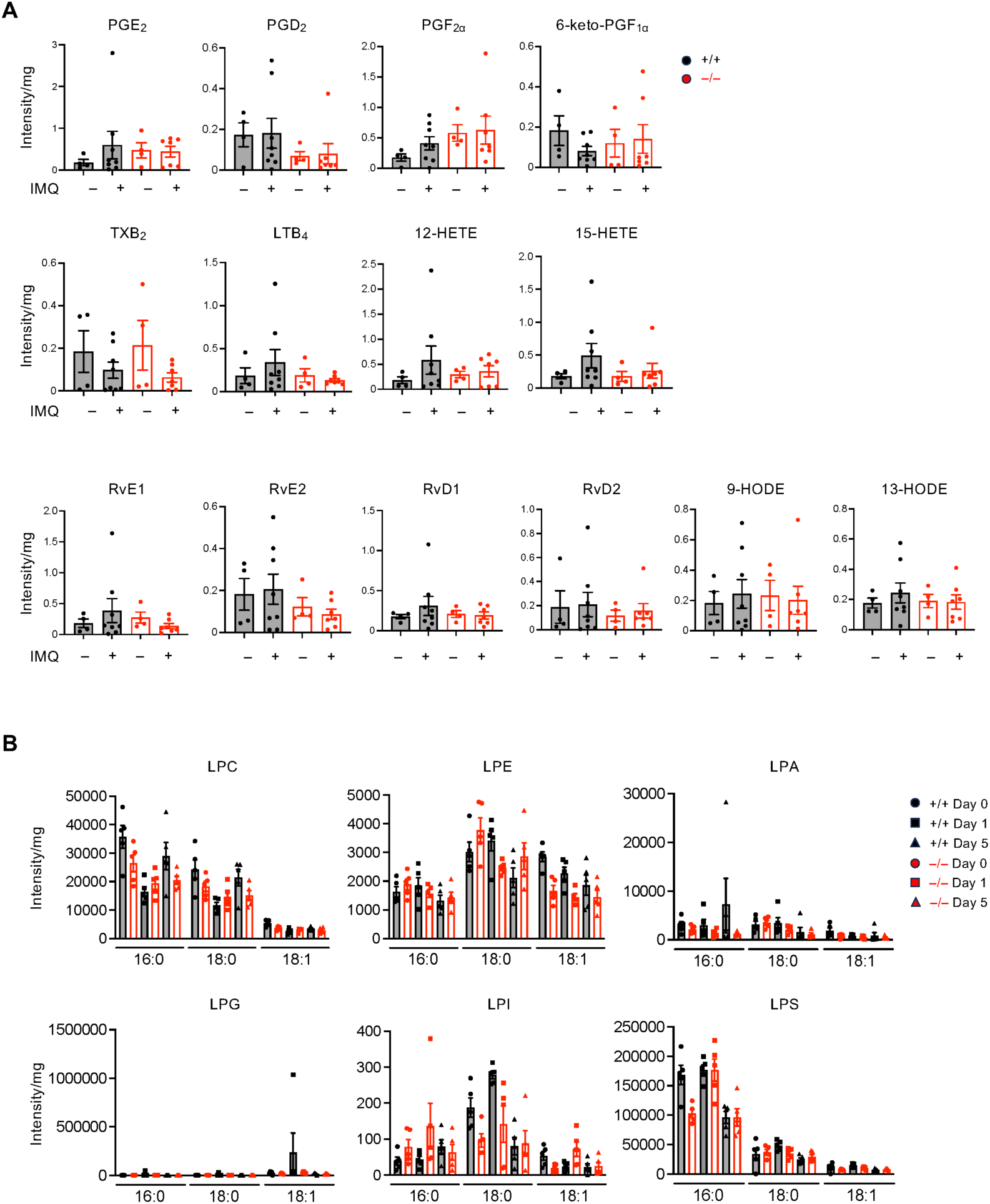
Lipidomics of the LNs and skin of *Pla2g12a*^+/+^ and *Pla2g12a*^-/-^ mice, related to Figure 5. (A) LC-ESI-MS/MS of PUFA metabolites in the LNs of *Pla2g12a*^+/+^ (+/+) and *Pla2g12a*^-/-^ (–/–) mice with (+) or without (–) IMQ treatment for 1 day. (B) LC-ESI-MS/MS of lysophospholipids in the skin of *Pla2g12a*^+/+^ and *Pla2g12a*^-/-^ mice treated for the indicated periods with IMQ. Values are mean ± s.e.m.. Results from one (A) or two (B) experiments are shown. Statistical analysis was performed using Brown-Forsythe and Welch ANOVA with Dunnett’s T3 multiple comparisons test (A) and two-way ANOVA with Sidak’s multiple comparisons test (B).

**Figure S7.**
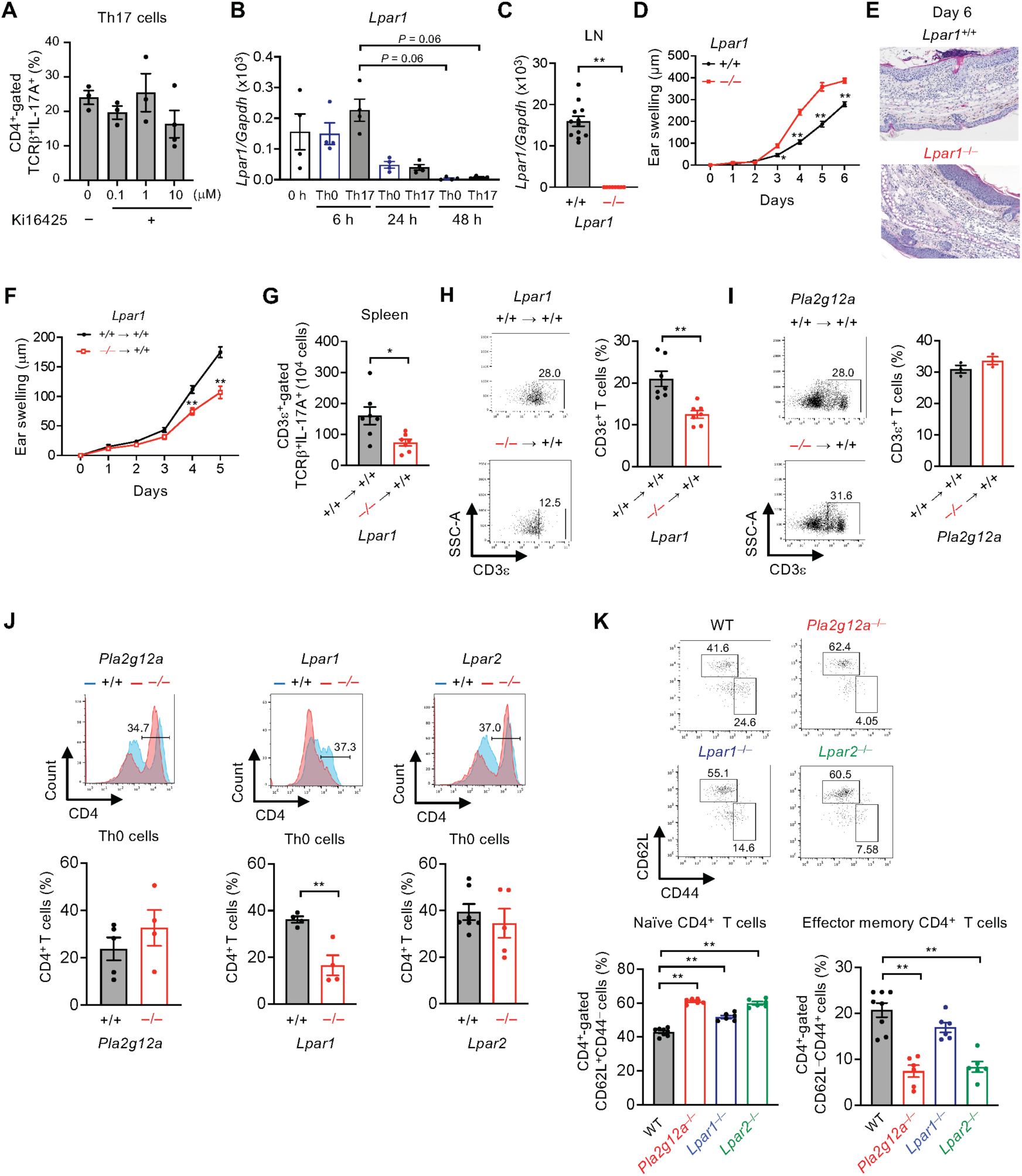
Analysis of LPA_1_- and LPA_2_-deficient mice, related to Figure 5. (A) FACS of Th17 cells after culture of naïve T cells for 3 days in the presence of the indicated concentrations of the LPA_1/3_ antagonist Ki16425. (B) qPCR of *Lpar1* in Th0 and Th17 cells after culture for the indicated periods. (C) qPCR of *Lpar1* in the LNs of *Lpar1*^+/+^ and *Lpar1*^-/-^ mice. (D, E) IMQ-induced ear swelling over 6 days (*n* = 8–12) (D) and ear histology on day 6 (E) of *Lpar1*^+/+^ and *Lpar1*^-/-^ mice. (F, G) IMQ-induced ear swelling over 6 days (*n* = 6) (F) and the number of splenic Th17 cells (G) of *Lpar1*^+/+^ mice that had been transferred with *Lpar1*^+/+^ or *Lpar1*^-/-^ BM cells. (H, I) FACS of splenic CDε+ T cells in *Lpar1*^+/+^ mice that had been transferred with *Lpar1*^+/+^ or *Lpar1*^-/-^ BM cells (H) or in *Pla2g12a*^+/+^ mice that had been transferred with *Pla2g12a*^+/+^ or *Pla2g12a*^-/-^ BM cells (I). Representative FACS profiles (*left*) and the proportion of CD3ε^+^ T cells (*right*) are shown. (J) FACS of CD4^+^ Th0 cells from *Pla2g12a*^-/-^, *Lpar1*^-/-^, and *Lpar2*^-/-^ mice in comparison with those from respective control mice. Representative FACS profiles (*upper*) and the proportion of CD4^+^ T cells (*lower*) are shown. (K) FACS of splenic CD4^+^-gated CD44^-^CD62L^+^ naïve T cells and CD44^+^CD62L^-^ effector memory T cells in *Pla2g12a*^-/-^, *Lpar1*^-/-^, and *Lpar2*^-/-^ mice in comparison with WT mice. Values are mean ± s.e.m.. Representative data of two experiments (B, J) and results of one experiment (A, C–I, K) are shown. Statistical analysis was performed using unpaired t test (C, I, J), Mann-Whitney U test (G, H), ordinary one-way ANOVA with Dunnett’s multiple comparisons test (A, K), Brown-Forsythe and Welch ANOVA with Dunnett’s T3 multiple comparisons test (B), and two-way ANOVA with Sidak’s multiple comparisons test (D, F). *, *P* < 0.05; **, *P* < 0.01.

**Figure S8.**
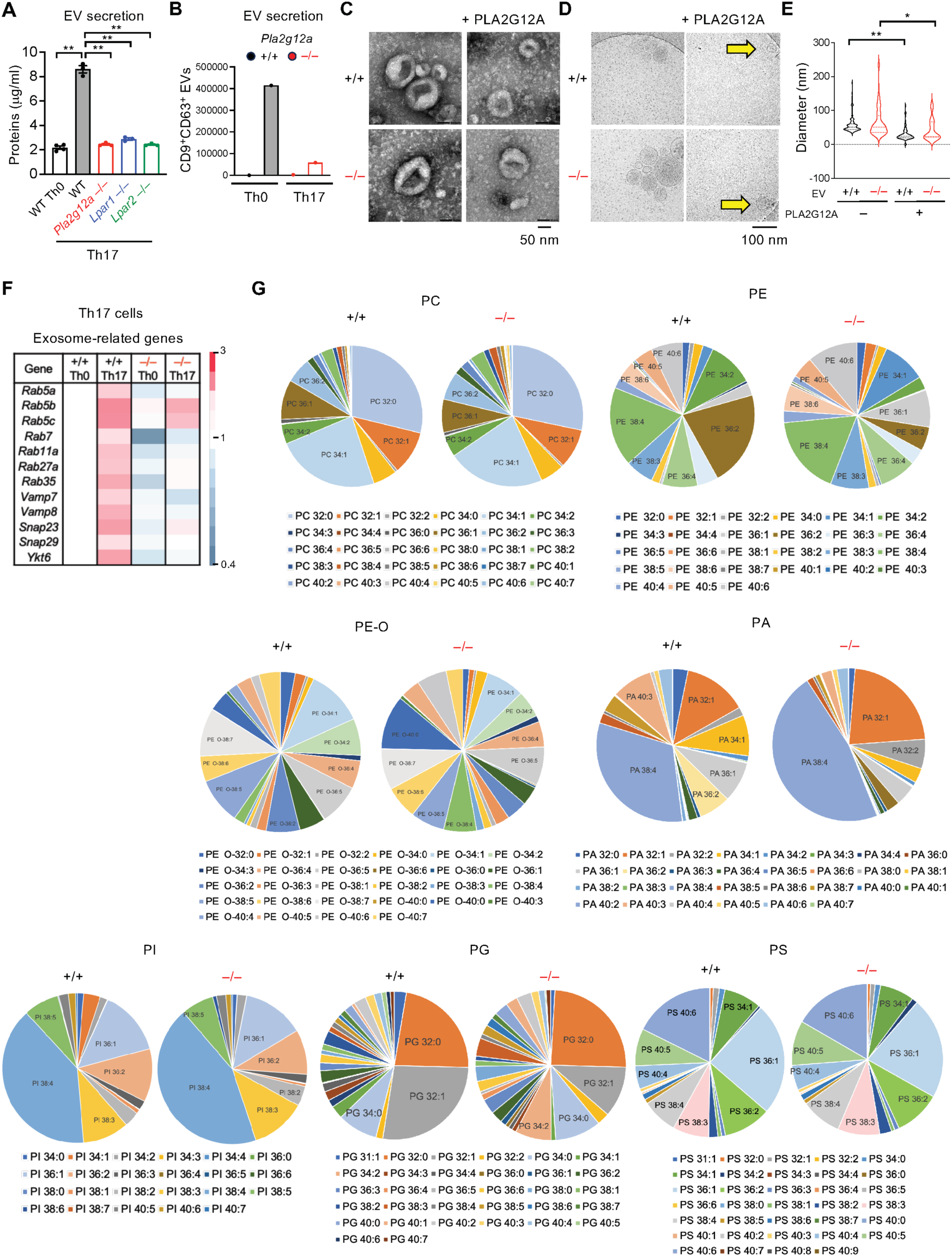
Secretion, size, and phospholipid composition of EVs from Th17 cells, related to Figure 6. (A, B) Secretion of EVs from Th0 and Th17 cells prepared from WT, *Pla2g12a*^-/-^, *Lpar1*^-/-^, and *Lpar2*^-/-^ mice after culture for 2 days, as evaluated by protein amounts (μg/ml of culture medium) (A) and counts of CD9^+^CD63^+^ EVs (B). In (B), EVs from six mice were pooled and subjected to the analysis. (C, D) TEM (C) and cryo-EM (D) of EVs secreted from *Pla2g12a*^+/+^ and *Pla2g12a*^-/-^ Th17 cells before or after treatment with PLA2G12A. (E) Diameters of Th17-derived EVs from *Pla2g12a*^+/+^ and *Pla2g12a*^-/-^ mice before or after treatment with PLA2G12A. (F) Heatmap of exosome-related genes in Th0 and Th17 cells from *Pla2g12a*^+/+^ and *Pla2g12a*^-/-^ mice, with expression levels (fold values) in *Pla2g12a*^+/+^-derived Th0 cells regarded as 1. (G) Phospholipid composition in Th17-derived EVs from *Pla2g12a*^+/+^ and *Pla2g12a*^-/-^ mice. Values are mean ± s.e.m.. Representative data of two experiments (C, G) and results of one experiment (A, B, D–F) are shown. Statistical analysis was performed using ordinary one-way ANOVA with Dunnett’s multiple comparisons test (A) and Kruskal-Wallis test with Dunn’s multiple comparisons test (E). *, *P* < 0.05; **, *P* < 0.01.

**Figure S9.**
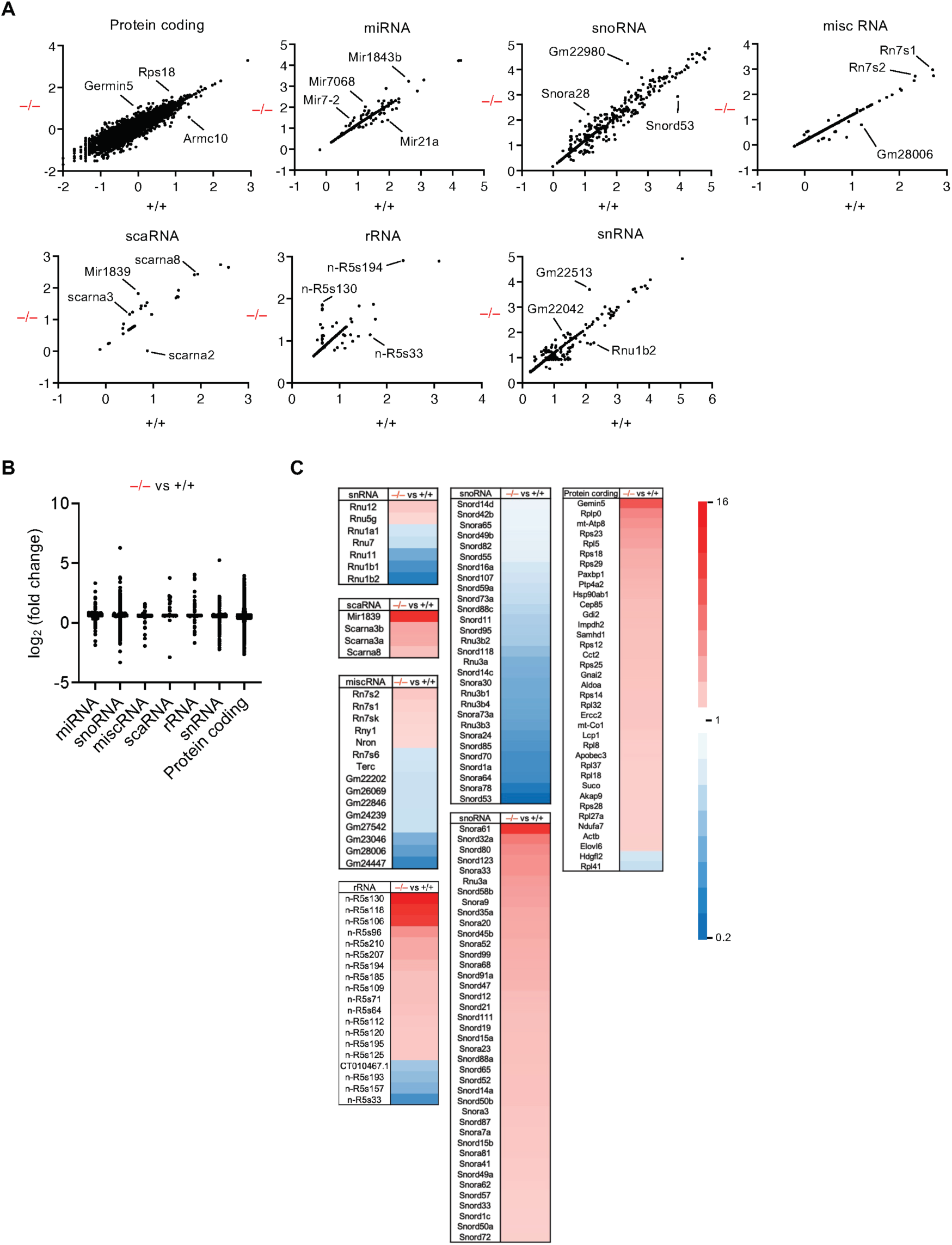
RNA-seq analysis of EVs from Th17 cells, related to Figure 6. (A) Scatter plots of various biotypes of RNA, including protein-coding (messenger) RNAs, miRNAs, small nucleolar RNAs (snoRNAs), small Cajal body-specific RNAs (scaRNAs), ribosomal RNAs (rRNAs), small nuclear RNAs (snRNAs), and miscellaneous RNAs (misc_RNAs) in Th17-derived EVs from *Pla2g12a*^+/+^ (+/+) and *Pla2g12a*^-/-^ (–/–) mice. (B) Abundances of individual RNA biotypes in Th17-derived EVs from *Pla2g12a*^-/-^ mice (–/– EVs) relative to those from *Pla2g12a*^+/+^ mice (+/+ EVs). (C) Heatmap of RNA biotypes whose levels were altered in Th17-derived EVs from *Pla2g12a*^-/-^ mice relative to those from *Pla2g12a*^+/+^ mice. Colors indicate fold changes in –/– EVs relative to +/+ EVs. One experimental data set, in which EVs from six mice were pooled, is shown.

**Figure S10.**
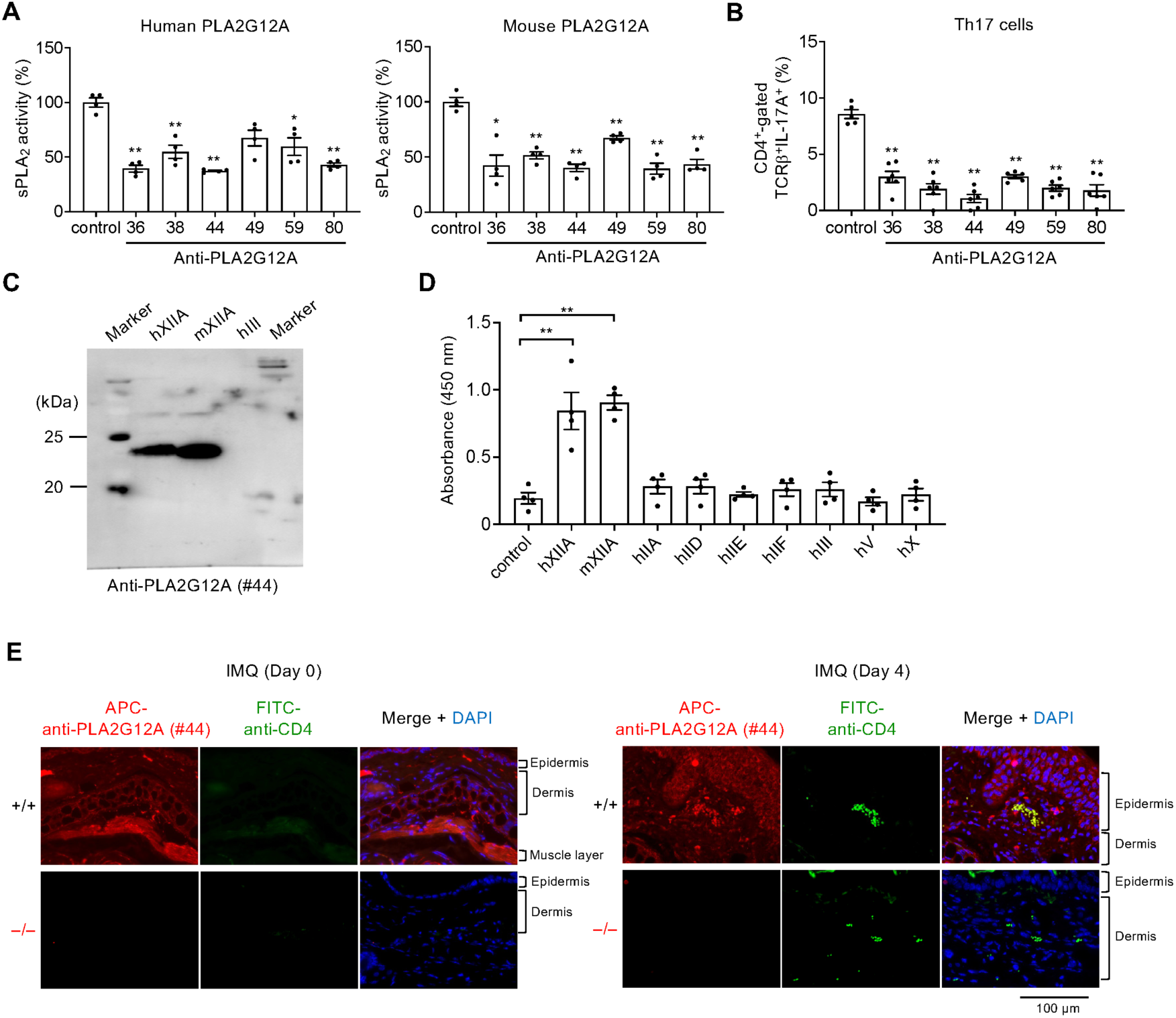
Establishment of anti-PLA2G12A monoclonal antibodies, related to Figure 7. (A) PLA_2_ enzyme assay using human and mouse PLA2G12A proteins after incubation with several anti-PLA2G12A antibody clones (100 ng/ml), with the activity in the presence of a control antibody being regarded as 1. *, *P* < 0.05; **, *P* < 0.01 *versus* control. (B) FACS of Th17 cells after culture of WT naïve T cells for 3 days with several anti-PLA2G12A antibody clones (100 ng/ml). *, *P* < 0.05; **, *P* < 0.01 *versus* control. (C) Immunoblotting of human and mouse PLA2G12As and human PLA2G3 (50 ng/lane for each) with an anti-PLA2G12A antibody (clone #44). (D) ELISA of various sPLA_2_s with an anti-PLA2G12A antibody (clone #44). hIIA, human PLA2G2A; hIID, human PLA2G2D; hIIE, human PLA2G2E; hIIF, human PLA2G2F; hIII, human PLA2G3; hV, human PLA2G5; hX, human PLA2G10; hXIIA, human PLA2G12A; mXIIA, mouse PLA2G12A. *, *P* < 0.05; **, *P* < 0.01. (E) Immunohistochemistry of mouse skin sections with or without IMQ treatment with an anti-PLA2G12A antibody (clone #44). Red, PLA2G12A; green, CD4^+^ T cells. Scale bar, 100 µm. Values are mean ± s.e.m.. Results of one experiment is shown (A–D). Statistical analysis was performed using Brown-Forsythe and Welch ANOVA with Dunnett’s T3 multiple comparisons test (A) and ordinary one-way ANOVA with Dunnett’s multiple comparisons test (B, D).

**Table S1.** TaqMan probe/primer sets for qPCR analysis.

| Gene Symbol | Assay No. | Source |
| --- | --- | --- |
| *Il6* | Mm00446190_m1 | Thermo Fisher Scientific |
| *Il17a* | Mm00439618_m1 | Thermo Fisher Scientific |
| *Il22* | Mm00444241_m1 | Thermo Fisher Scientific |
| *Il17f* | Mm.PT.58.9739903.g | Integrated DNA Technologies |
| *Rorc* | Mm.PT.58.23464726.g | Integrated DNA Technologies |
| *Il23ap19* | Mm01160011_g1 | Thermo Fisher Scientific |
| *Pla2g12a* | Mm01316982_m1 | Thermo Fisher Scientific |
| *Krt10* | Mm03009921_m1 | Thermo Fisher Scientific |
| *Flg* | Mm01716522_m1 | Thermo Fisher Scientific |
| *Lpar1/Edg2* | Mm00439145_m1 | Thermo Fisher Scientific |
| *Lpar2/Edg4* | Mm00469562_m1 | Thermo Fisher Scientific |
| *Lpar3/Edg7* | Mm00469694_m1 | Thermo Fisher Scientific |
| *Lpar4* | Mm02620784_s1 | Thermo Fisher Scientific |
| *Lpar5* | Mm02621109_s1 | Thermo Fisher Scientific |
| *Lpar6* | Mm00613058_s1 | Thermo Fisher Scientific |
| *Gapdh* | 4352339E | Thermo Fisher Scientific |

**Table S2.** Cell surface markers for FACS analysis.

| antibody | label | dilution | clone | Source |
| --- | --- | --- | --- | --- |
| CD45 | BV421 | 1:1,000 | 30-F11 | BioLegend |
| CD3ε | APC/Fire 750 | 1:500 | 145-2C11 | eBiosciences |
| CD4 | APC | 1:500 | GK1.5, RM4.5 | BioLegend |
| CD44 | BV480 | 1:500 | IM7 | BD |
| CD62L | FITC | 1:500 | MEL-14 | BioLegend |
| TCRβ | FITC<br>APC Cyanine7 | 1:500 | H57-597 | BioLegend |
| TCRγδ | FITC<br>APC Cyanine7 | 1:500 | GL3 | BioLegend |

BV (brilliant violet); APC (allophycocyanin); PE (phycoerythrin); FITC (fluorescein isothiocyanate)

